# Orthogonal CRISPR screens and human liver chimeric mice identify hepatitis B virus host factors

**DOI:** 10.64898/2026.07.23.740386

**Authors:** Catherine A Freije, Georgios Dangas, Antonis Athanasiadis, Evgenia Moschogianni, Madeleine K Sanders, Allan Henrique Depieri Cataneo, Chloe K Boehm, Maria Bousali, Dar-Yin Li, Sooyoung Lee, Guillaume Cornelis, Gary Lo, Leah B Soriaga, Amalio Telenti, Julia di Iulio, Ana L P Mosimann, Juliano Bordignon, Kirin Karver, Lukas Fu, Kenneth C Levenson, Chenhui Zou, Yichen Zhou, Corrine Quirk, Leon L Seifert, Xupeng Hong, Anushka Rajesh, Yingpu Yu, Andreas S Puschnik, Florian A Lempp, Herbert W Virgin, Seungmin Hwang, William M Schneider, Ype P de Jong, Charles M Rice, Eleftherios Michailidis

**Author notes:** These authors contributed equally: Catherine A Freije and Georgios Dangas. These authors contributed equally: Antonis Athanasiadis and Evgenia Moschogianni. These authors contributed equally: Charles M Rice and Eleftherios Michailidis. Instituto Carlos Chagas, Fiocruz Paraná, Fundação Oswaldo Cruz, Fiocruz, Curitiba, PR, Brazil. The Gates Foundation, Seattle, WA 98109, USA.

## Abstract

Hepatitis B virus (HBV) chronically infects approximately 250 million people worldwide, and reliable curative therapies are lacking. A broader understanding of viral-host interactions could accelerate efforts to find new host-centric therapeutic targets. However, inefficient cell culture systems and limited replication markers compatible with pooled screening have precluded the widespread use of genetic perturbation screens. Here, we performed the first pooled, genome-wide CRISPR-Cas9 knockout (KO) screen with authentic HBV infection and integrated these results with two orthogonal pooled screens to identify host factors. We selected 72 genes for a multi-step assessment that included arrayed validation assays using both HBV infection and pgRNA transfection. We then independently tested thirteen genes using high-efficiency bulk KO experiments to guide further investigations of both antiviral and proviral factors. In both KO and siRNA-mediated knockdown experiments, depletion of the top antiviral factor, *EXOC1*, enhanced multiple HBV replication markers, and transcriptomic analysis revealed activation of hypoxia- and HIF-1 gene signatures. Three proviral factors, *IRF2*, *WDR48*, and *ZCCHC14*, were investigated *in vivo* using a human liver chimeric mouse model, which demonstrated that *ZCCHC14* KO greatly reduced HBV replication and spread. Together, these complementary *in vitro* and *in vivo* platforms expand the catalog of HBV host factors and provide a scalable framework for host target discovery.

## Introduction

Despite effective vaccines, HBV remains a global health concern with approximately 250 million chronic infections worldwide and no reliable curative therapies^1^. Currently approved antiviral therapies include interferon alpha and nucleos(t)ide analogs, but they rarely cure infection and incompletely mitigate long-term morbidity and mortality^2,3^. The persistence of a stable, episomal viral genome, i.e., the covalently closed circular DNA (cccDNA), is key to establishing chronic infection. Therefore, many therapeutic efforts have been directed towards cccDNA elimination or silencing, as well as strategies to promote the reduction or loss of circulating HBV surface antigen (HBsAg), a feature of a functional cure^2,4^.

HBV is a small (3.2 kb), partially double-stranded DNA virus that exclusively infects human hepatocytes and engages with a broad network of host factors^5^. Entry begins when the viral surface protein engages heparan sulfate proteoglycans^6^ followed by specific binding to the entry receptor, sodium taurocholate co-transporting polypeptide (NTCP)^7^. Once virions enter cells, nucleocapsids containing the partially double-stranded relaxed circular DNA (rcDNA) traffic to the nucleus, relying on endosomal-associated proteins Rab5 and Rab7^8^ and microtubules^9^. Then, DNA repair machinery converts rcDNA to fully double-stranded cccDNA, a process that involves removal of the associated HBV polymerase^10^ and small RNA primer^11^ followed by filling in the DNA gap and ligation^12,13^. Once chromatinized, cccDNA serves as the template for viral transcription, generating five primary viral mRNAs. Transcription is regulated by hepatocyte-specific transcription factors such as HNF4α, RXRα, and PPARα^14^. Despite much effort, our understanding of the host factors that either promote or inhibit these stages of HBV replication in human hepatocytes remains incomplete.

Genetic perturbation screens using small-interfering RNAs (siRNAs) or CRISPR systems are powerful tools for identifying host factors involved in various biological processes. Yet, pooled screens rely on selectable or sortable phenotypes, and viral spread or virus-mediated cell death can provide strong selection pressures that enable robust enrichment of host-factor perturbations^15^. Because of HBV’s noncytopathic nature and low infectivity in cell culture^16^, prior pooled genetic screens for HBV have been limited to those that employed host factor overexpression^17^, investigated HBV markers expressed from an integrated intermediate^18^, or, most recently, used a reporter virus^19^. These constraints have also motivated the use of arrayed siRNA screens, in which genes are tested individually^20^. These prior screen datasets provide valuable starting points for more detailed investigations of putative HBV host factors. However, to our knowledge, no pooled knockout (KO) screen has been performed using an authentic/non-reporter HBV. Further, establishing a set of assays to determine which of the hundreds of potential HBV host factors identified in such screens have robust effects across *in vitro* and *in vivo* HBV replication models is critical to elucidating HBV’s host factor network.

Here, we performed the first pooled, genome-wide CRISPR-Cas9 KO screen using *de novo* HBV infection, identifying hundreds of putative HBV host factors. We then established an orthogonal arrayed KO approach to validate 72 of the screen-identified hits in multiple hepatocyte HBV systems. We integrated the results from the arrayed validation experiments with a targeted, literature-informed analysis focusing on genes hypothesized to play roles in intermediate life-cycle steps, resulting in the selection of 13 genes for further study. One antiviral factor, *EXOC1*, was studied across multiple HBV systems. In addition, three proviral targets (*IRF2*, *WDR48*, and *ZCCHC14*) were selected for *in vivo* evaluation in a human liver chimeric mouse model that supports robust HBV infection and spread.

## Results

### Pooled CRISPR-Cas9 KO screens to identify HBV host factors

To identify host factors involved in HBV infection, we performed a genome-wide CRISPR KO screen in HepG2-NTCP cells using antibody-stained HBsAg as a marker for productive infection (Fig. 1a). The pooled KO population of HepG2-NTCP was generated by transducing cells with the Brunello lentiviral library that expresses both Cas9 and single guide RNAs (sgRNAs)^21^. This library targets 19,114 human genes with four sgRNAs per gene and includes 1,000 non-targeting (NT) sgRNAs. Cells were then infected with HBV genotype D derived from HBV-producing HepAD38 cells, followed by intracellular HBsAg staining and fluorescence-activated cell sorting (FACS). The relative abundance of sgRNAs was then assessed by sequencing the genomic DNA of the sorted populations.

**Fig. 1.**
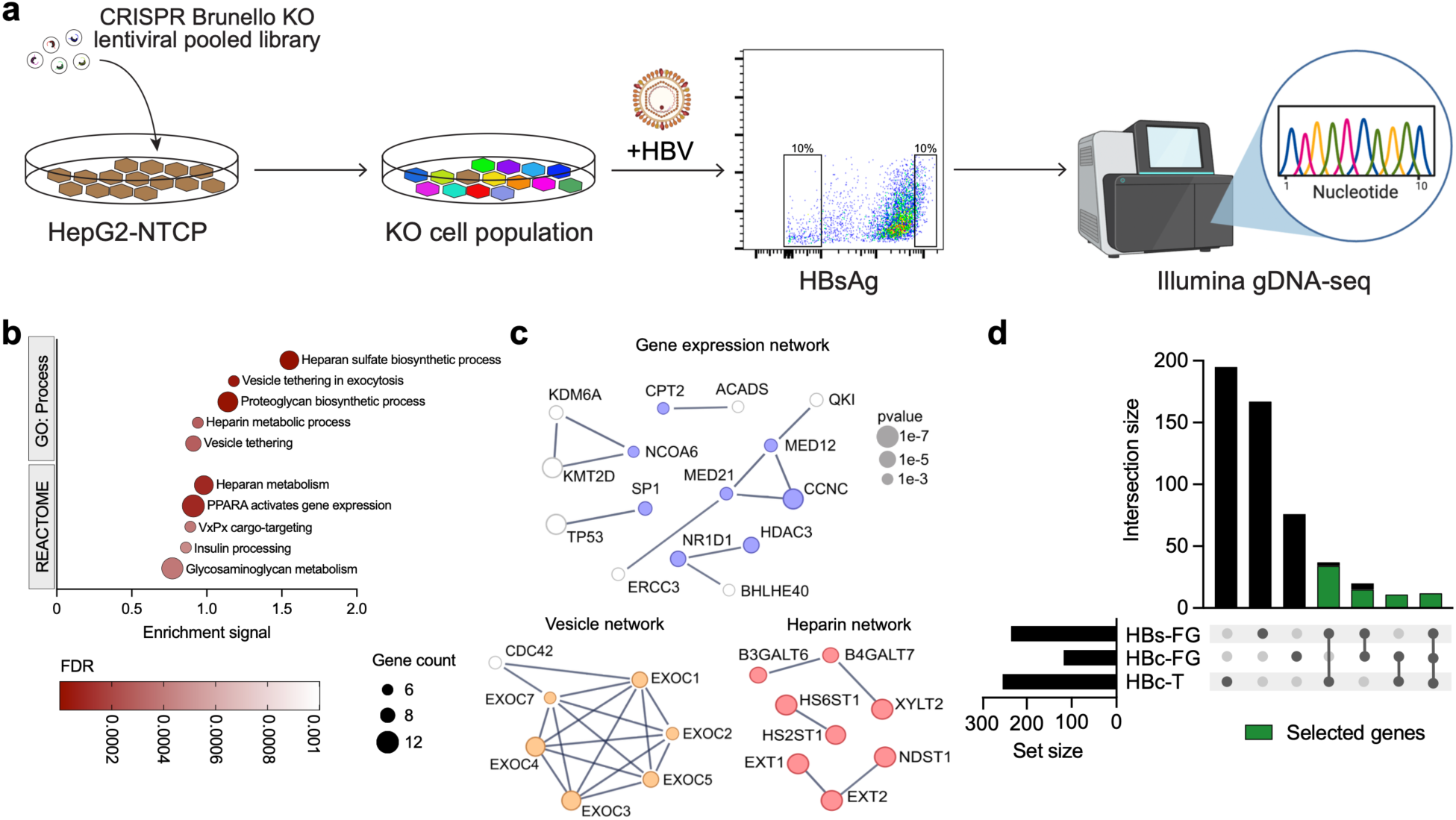
Genome-wide CRISPR-Cas9 KO screen for identification of HBV host dependency factors. **a** Schematic of the FACS-based, genome-wide CRISPR-Cas9 KO screen with HBV infection. Created with BioRender.com. **b** Gene set enrichment analysis of the significant genes identified in the KO screen (significant genes in database = 229). **c** Functional network of the significant genes contained within the enriched gene sets using STRINGdb^48^. Only the highest-confidence (score ≥ 0.9) interactions are included. First-degree connections to other significant genes but not in gene sets are shown in white. Symbol size scaled relative to *P* in the screen. **d** UpSet plot of the overlapping genes identified between the three orthogonal HBV infection screens performed. Selected hits for follow-up studies are shown in green to indicate that the selection required the gene to pass significance thresholds in at least two screens. HBs-FG, HBsAg full-genome screen hits; HBc-FG, HBcAg full-genome screen hits; HBc-T, HBcAg targeted screen hits.

By measuring sgRNA abundance in infected and sorted HepG2-NTCP cells relative to the unsorted population, we identified a set of putative host factors that modulate HBsAg levels. Using a p-value threshold of 1e-3, 236 genes were identified as potential HBV host factors (Supplementary Table 1). Importantly, known HBV host factors including hepatocyte nuclear factor 4 alpha (*HNF4A*)^14^, DNA polymerase κ (*POLK*)^12,13^, and zinc finger CCHC-type containing 14 (*ZCCHC14*)^18^ were among the hits with the applied significance threshold.

To further characterize the screen hits, we performed gene set enrichment analysis. Assessment of multiple gene set databases uncovered a significant enrichment of pathways related to heparan sulfate biosynthesis, vesicle tethering, and gene expression (Fig. 1b, Supplementary Table 2). To visualize the functional interactions of genes that passed the screen significance threshold, we generated a complete interactome map of the selected hits (229/236 genes mapped to the database) and highlighted the genes that were identified in the enriched gene set and pathway analysis (Fig. 1c, Supplementary Fig. 1a). The identification of heparan sulfate biosynthesis pathways could be related to known, low-affinity interactions between HBsAg and heparan sulfate proteoglycans, thereby affecting viral entry^6,22^ or impacting intracellular or surface-associated HBsAg because HBsAg was used as the screen readout. Vesicle tethering gene sets were significantly enriched, largely due to the inclusion of six of the eight members of the exocyst complex in our list of screen hits. The exocyst complex has previously been reported to play a role in SARS-CoV-2^23^, HIV^24^, and herpesvirus replication^25^. However, its role in HBV infection has not been established. Finally, the PPARA gene expression set included two members of the mediator complex, *MED12* and *MED21*, a regulator of RNA polymerase II-based transcription^26,27^.

We compared the selected genes identified in our screen with those from two prior screens^18,20^. These two screens differed from the screen performed here, as one used an integrated HBV replication model^18^ and the other used an arrayed siRNA approach^20^. Unsurprisingly, given the technical differences among the screens and the signal-to-noise of HBV replication markers, the overlap among the three screens was small (Supplementary Fig. 1b, Supplementary Table 3). Only two host factors were shared across all three screens: *CDKN1A*, encoding p21, a known cell cycle regulator often less mutated in virus-associated cancers^28,29^, and *KHDRBS1*, an RNA-binding protein shown to stabilize hepatitis delta virus RNAs^30^.

For the pair-wise comparison between our screen and the screen with the integrated HBV replication model^18^, the overlap between the two pooled CRISPR-Cas9 KO screens was more than expected by chance (Fisher’s exact test, odds-ratio: 6.2, *P =* 2.6e-7) despite being performed in different cell lines and HBV replication models. This overlap motivated us to conduct two additional, orthogonal CRISPR KO screens to prioritize targets for downstream validation.

We performed two orthogonal screens and established a set of priority targets for follow-up studies. The first was performed using the above-described Brunello lentiviral library; HBV infection was then assessed by intracellular staining for HBV core antigen (HBcAg), followed by FACS sorting. The second was a targeted screen, in which the top 960 genes ranked by p-value from the genome-wide HBsAg screen were used to generate a custom lentiviral library and screened using the HBcAg readout. Analysis of the overlapping screen hits led to the selection of 72 genes for further study (Fig. 1d, Supplementary Fig. 1c). Collectively, these complementary screening datasets identified a robust set of candidate HBV host factors and laid the foundation for subsequent arrayed validation studies.

### Arrayed CRISPR-Cas9 KO validation identifies candidate host dependency and restriction factors during HBV infection

To validate the 72 screen-selected candidate HBV host factors, we performed an arrayed CRISPR-Cas9 KO validation assay in HepG2-NTCP cells (Fig. 2a). HepG2-NTCP cells were transfected with a pool of sgRNAs complexed with recombinant Cas9 nuclease in a 96-well arrayed format (Supplementary Table 4). To allow time for KO and protein turnover, cells were cultured for 5 days prior to HBV infection, and viral markers were quantified 7 days post-infection using both extracellular and intracellular viral readouts.

**Fig. 2.**
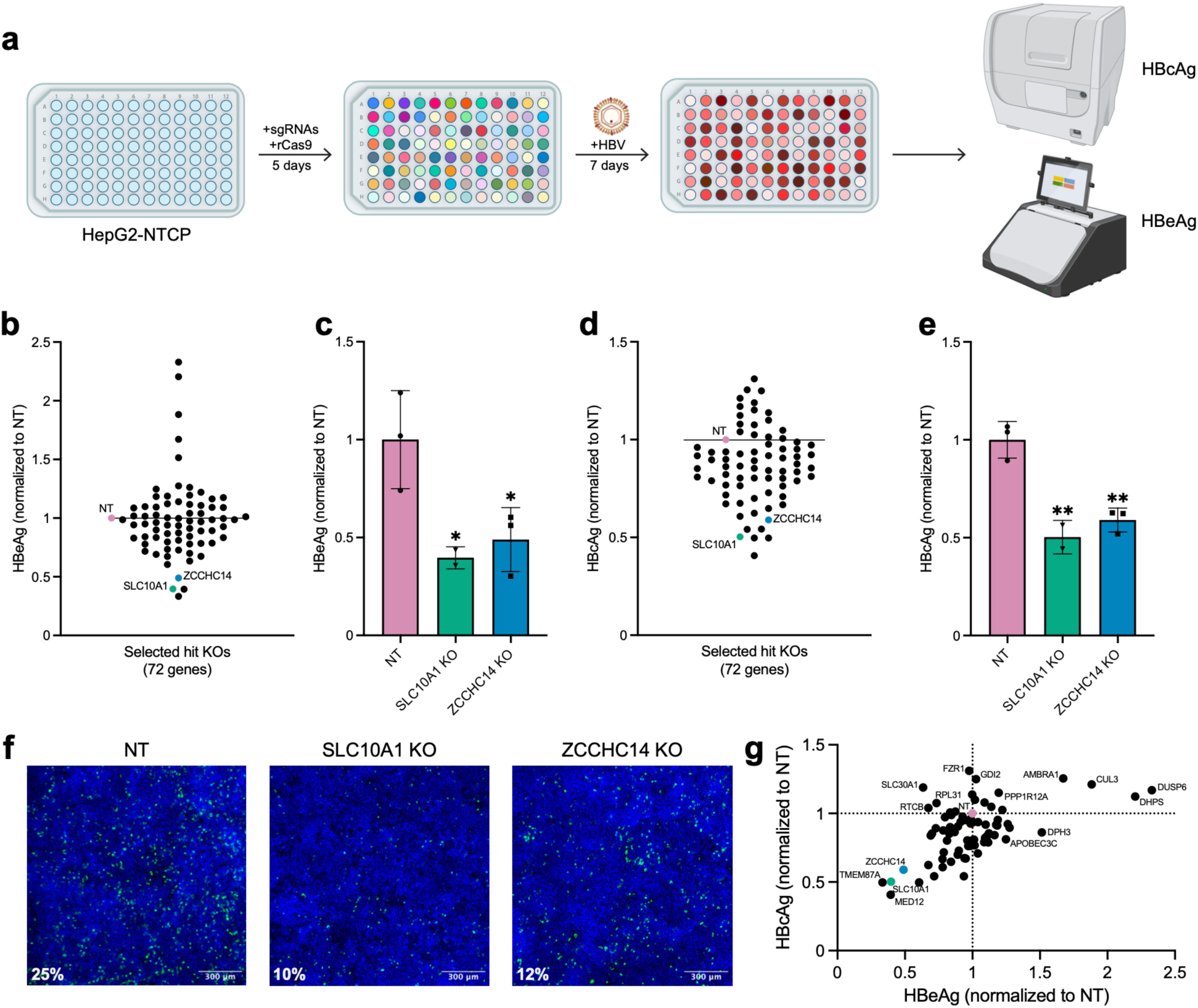
Arrayed CRISPR-Cas9 KO validation of candidate host factors identified in the genome-wide HBV screen using an HBV infection model. **a** Schematic overview of the HBV infection-based validation workflow. HepG2-NTCP cells were transfected in an arrayed 96-well format with recombinant Cas9 nuclease and sgRNAs targeting the 72 selected genes. Five days post-transfection (dpt), cells were infected with HBV and maintained for an additional 7 days prior to analysis. HBV infection was quantified by measuring secreted HBeAg levels in the supernatant and intracellular HBcAg levels by immunofluorescence staining. Red shading in the assay plate represents relative levels of HBV infection. Created with BioRender.com. **b** Quantification of HBeAg levels in cell culture supernatant 7 days post-infection (dpi) in the generated KO cells. HBeAg levels were normalized to cell number as measured by Hoechst staining and subsequently normalized to the non-targeting sgRNA control (NT). Each symbol represents the mean of 3 replicates for a given gene KO. **c** Measurement of secreted HBeAg levels for *SLC10A1* and *ZCCHC14*. **d** Quantification of intracellular HBcAg staining following knockout of the 72 selected candidate genes. HBcAg signal was normalized to cell number and subsequently normalized to NT controls. Each symbol represents the mean of 3 replicates for a given gene knockout. **e** HBcAg staining measurements for *SLC10A1* and *ZCCHC14*. **f** Representative immunofluorescence images of HBcAg staining in NT (25% positive), *SLC10A1* KO (10% positive), and *ZCCHC14* KO (12% positive) cells. Scale bars, 300 μm. **g** Correlation plot comparing normalized HBeAg and HBcAg readouts across the 72 candidate gene KOs. Strongly correlated and anticorrelated proviral host dependency factors and antiviral host restriction factors are labeled. Data in **c** and **e** are presented as mean ± S.D. of three replicates. Statistical significance was assessed by ordinary one-way ANOVA followed by Dunnett’s multiple comparisons test using NT as the reference group. \**P* < 0.05; \*\**P* < 0.01.

We first quantified secreted HBV e antigen (HBeAg) levels in the culture supernatant to assess productive HBV infection. HBeAg levels were normalized to cell number and subsequently normalized to the NT control sgRNA (Fig. 2b). As expected, KO of *SLC10A1*, which encodes the HBV entry receptor NTCP, resulted in a marked reduction in HBeAg levels (Fig. 2c). Similarly, KO of *ZCCHC14* reduced HBeAg production, consistent with its previously reported role in HBV replication^18^.

To assess intracellular markers of HBV infection, we quantified HBcAg levels by immunofluorescence staining (Fig. 2d). Consistent with the HBeAg measurements, KO of both *SLC10A1* and *ZCCHC14* significantly reduced HBcAg expression relative to NT controls (Fig. 2e). Representative immunofluorescence images further confirmed the reduction of HBcAg-positive cells following disruption of these positive control genes (Fig. 2f).

Comparison of extracellular HBeAg and intracellular HBcAg readouts across all candidate genes demonstrated that many genes showed a positive correlation (Fig. 2g, Pearson r = 0.54, P < 0.0001). Several candidate genes reproducibly suppressed or enhanced HBV infection upon KO, identifying potential proviral host dependency factors (33 out of 72 candidates) and antiviral host restriction factors (11 out of 72 candidates) for further characterization. Overall, the concordance between extracellular and intracellular infection readouts validated the arrayed screening approach and identified a subset of candidate proviral and antiviral host factors for further study.

### Initiating HBV replication with pgRNA distinguishes entry-dependent and post-entry host factors

To determine which of the screen-identified host factors regulate post-entry stages of the HBV replication cycle, we employed a method that bypasses viral entry (Fig. 3a)^31,32^. During HBV’s life cycle, a full-genome length RNA intermediate, the pregenomic RNA (pgRNA), is transcribed from cccDNA. Transfection of pgRNA can initiate HBV genome replication, leading to detectable cccDNA and viral transcription products including HBsAg^31,32^. Therefore, we reasoned that this approach would specifically identify host factors that regulate post-entry processes.

**Fig. 3.**
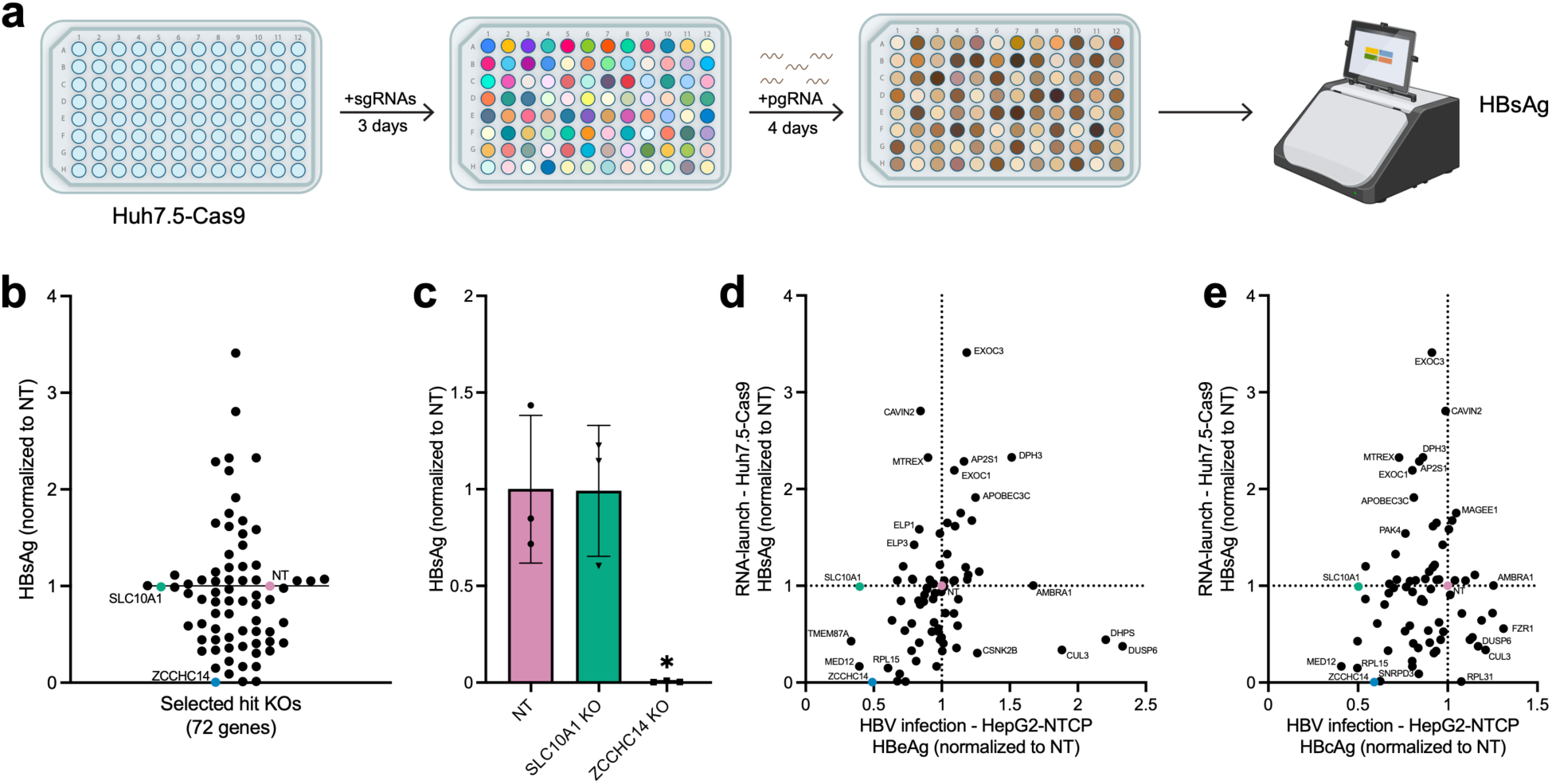
Arrayed CRISPR-Cas9 KO validation identifies host factors regulating post-entry stages of the HBV life cycle. **a** Schematic overview of the arrayed CRISPR-Cas9 KO validation workflow using the HBV pgRNA launch system. Huh7.5-Cas9 cells were transfected in an arrayed 96-well format with sgRNAs targeting the selected 72 genes. Three dpt of sgRNAs, cells were transfected with HBV pgRNA, and the supernatant was harvested 4 dpt of pgRNA for quantification of secreted HBsAg. Brown shading in the assay plate represents relative levels of HBV replication. Created with BioRender.com. **b** Quantification of secreted HBsAg levels following knockout of the 72 selected genes. HBsAg levels were normalized to cell number and subsequently normalized to the NT sgRNA. Each symbol represents the mean of 3 replicates for a given gene knockout. **c** Measurement of secreted HBsAg levels for *SLC10A1* and *ZCCHC14*. **d,e** Correlation of normalized secreted HBsAg levels following pgRNA transfection (y-axis) with HBV infection readouts from Fig. 2, including normalized HBeAg levels (d) and normalized HBcAg levels (e). Host dependency factors and antiviral host restriction factors showing concordant or discordant effects across the two validation systems are labeled. Data in **c** are presented as mean ± S.D. of three replicates. Statistical significance was assessed by ordinary one-way ANOVA followed by Dunnett’s multiple comparisons test using NT as the reference group. \**P* < 0.05.

Using the same sgRNA pools tested in the HepG2-NTCP arrayed infections (Supplementary Table 4), we generated individual gene KOs in Huh7.5 cells stably expressing Cas9 (Huh7.5-Cas9) and then transfected them with HBV pgRNA. HBV replication was assessed by quantifying HBsAg levels in the supernatant normalized to cell number (Fig. 3b, Supplementary Table 5). HBsAg was used to measure replication efficiency, as HBc abundance is biased by the core protein production from input pgRNA, whereas HBsAg can originate only from *de novo-*established HBV DNA.

As expected for an assay that bypasses viral entry, KO of *SLC10A1* did not reduce HBsAg levels following pgRNA transfection (Fig. 3c). In contrast, *ZCCHC14* KO markedly reduced HBsAg production.

Comparing results across arrayed screening datasets revealed both shared and distinct host factor dependencies between the two systems (Fig. 3d,e). Several candidate genes exhibited concordant phenotypes in both assays (e.g., *EXOC3*, *DPH3*, *AP2S1*, *EXOC1*, *APOBEC3C, MED12, TMEM87A, RPL15*), suggesting functions in post-entry stages of the HBV life cycle, whereas others displayed divergent phenotypes consistent with roles in viral entry or early post-entry processes (e.g., *AMBRA1*, *RABIF*, *SLC10A1*, *CUL3*, *DHPS*, *DUSP6*, *CSNK2B, FOXK1*). Together, these results distinguished host factors acting at viral entry from those regulating post-entry stages of the HBV life cycle and provided an additional layer of functional prioritization for downstream studies.

### Integrated analysis of the screen hits

Hepatocyte cell lines are a convenient source of cells for both pooled and arrayed screens, but these cells may not always recapitulate the cellular state of primary hepatocytes and, consequently, HBV-host factor dependencies^33^. Therefore, we did not exclusively rely on the arrayed screening results to select a smaller set of genes for further studies. The 72 screen-identified genes were additionally subject to an in-depth, literature-informed analysis (Supplementary Fig. 2a). After excluding genes hypothesized to be involved in entry, trafficking, mitochondria, and translation, 28 genes were assessed according to gene function, toxicity, and potential drug availability as a measure of potential translational impact (Supplementary Table 5).

The 28 genes independently selected from the literature-informed analysis were then combined with the list of genes validated in at least one of the arrayed screens (38 genes). This combined list included 53 genes, as 13 overlapped between the two analyses (Supplementary Table 5). We then selected 13 of the 53 genes incorporating both the arrayed screening results and assessed metrics defined above, for further study.

To validate these candidates in an independent KO format, we generated stable HepG2-NTCP KO cell lines using a lentiviral CRISPR-Cas9 approach. For each target gene, two new sgRNAs were designed and introduced into the LentiCRISPRv2 plasmids (one guide per plasmid), which co-express the Cas9 nuclease and puromycin resistance^34^. Lentiviral stocks were generated for each sgRNA and transduced at equal ratios into HepG2-NTCP cells. Transduced cells were selected with puromycin to enrich for the Cas9-sgRNA-expressing cell population and then infected with HBV. Intracellular HBcAg and secreted HBeAg were measured 7 days post-infection. To establish this approach, additional control lentiviruses were included alongside NT and *SLC10A1*: *HIRA*, *DDB1*, and *PCNA*. Of the 13 selected genes, *ZCCHC14*, *IRF2*, *WDR48*, and *FOXK1* KOs reduced both HBeAg and HBcAg-positivity, whereas *EXOC1* KO showed the greatest increases in both replication markers (Fig. 4a-d). These results identified *EXOC1* as the strongest antiviral candidate and *ZCCHC14*, *IRF2*, *WDR48*, and *FOXK1* as candidate proviral host factors for further study.

**Fig. 4.**
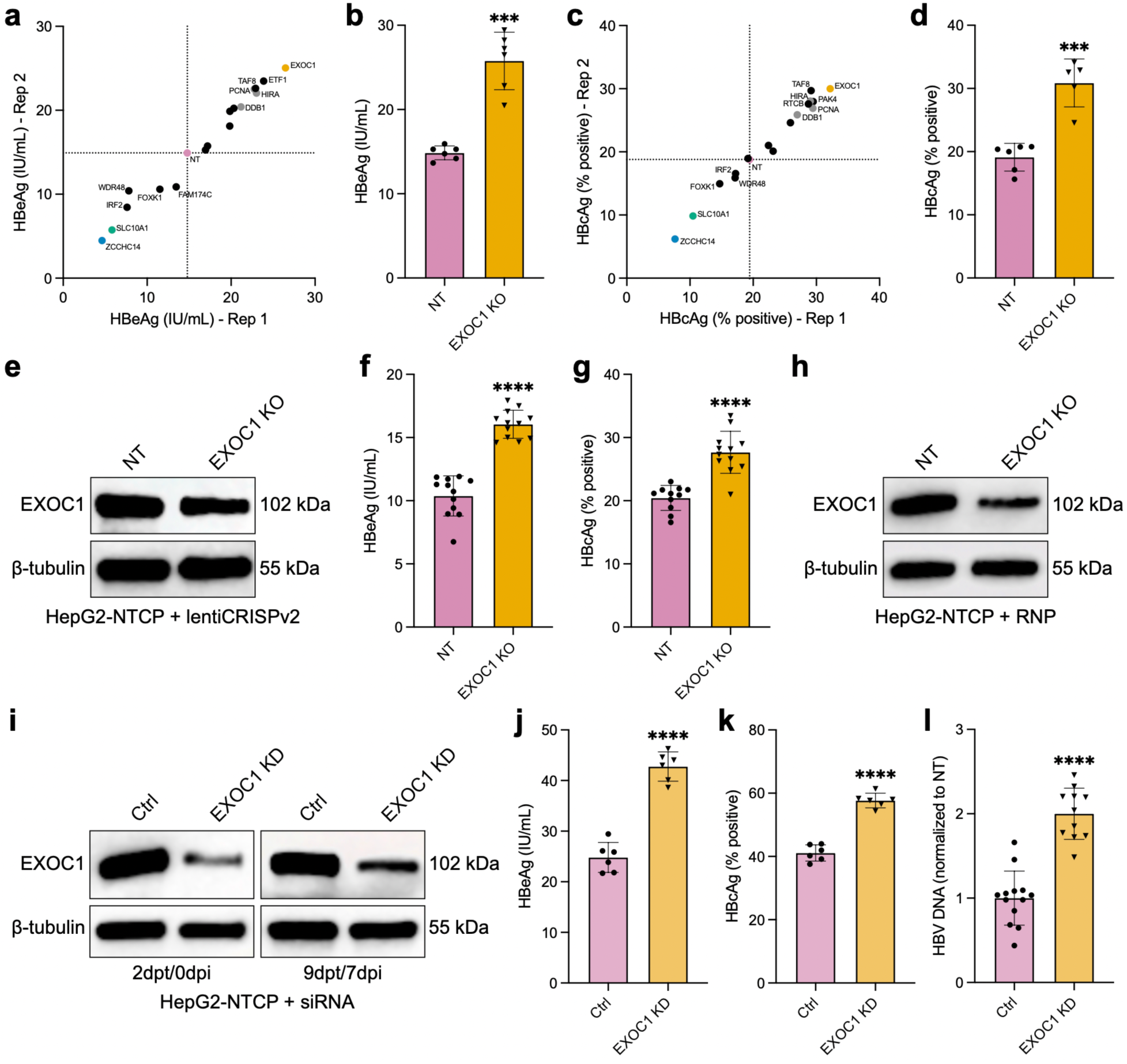
Orthogonal validation identifies EXOC1 as a candidate host factor restricting HBV infection. **a** HBcAg immunofluorescence-based validation of prioritized candidate genes: 13 selected gene KOs and controls in stable HepG2-NTCP knockout cell lines generated using a lentiviral CRISPR-Cas9 approach. Transduced cells were infected with HBV, and HBcAg-positive cells were quantified 7 dpi. Data are shown as the percentage of HBcAg-positive cells across two independent biological replicates of HBV infections. Positive control knockouts targeting *SLC10A1* and *ZCCHC14* reduced HBcAg positivity, whereas *EXOC1* KO increased HBcAg positivity relative to non-targeting (NT) controls. Each symbol represents the mean of three technical replicates from each independent HBV infection experiment (n = 6). NT: pink; *SLC10A1 KO:* green; *ZCCHC14* KO: blue; *DDB1*/*PCNA*/*HIRA KOs:* grey. **b** Replicate measurements of HBcAg-positive cells in NT and *EXOC1* KO HepG2-NTCP cells from results shown in **a**. **c** HBeAg-based validation of the prioritized candidate genes in stable HepG2-NTCP knockout cell lines. Secreted HBeAg levels were quantified in culture supernatants 7 dpi across two independent biological replicates of HBV infections. Each symbol represents the mean of three technical replicates from each independent HBV infection experiment (n = 6). **d** Replicate measurements of supernatant HBeAg levels in NT and *EXOC1* KO HepG2-NTCP cells from the validation experiment shown in panel **c**. **e** Western blot analysis of EXOC1 protein levels in NT and *EXOC1* KO HepG2-NTCP cells generated using the lentiviral CRISPR-Cas9 approach. β-tubulin was used as a loading control. **f, g** Orthogonal validation of *EXOC1* KO in HepG2-NTCP cells using recombinant Cas9 and synthetic sgRNAs. Cells were infected with HBV, and infection was quantified by HBcAg immunofluorescence staining **(f)** and supernatant HBeAg levels **(g)** 7 dpi. **h** Western blot analysis of EXOC1 protein levels in NT and *EXOC1* KO HepG2-NTCP cells generated using recombinant Cas9 and synthetic sgRNAs. β-tubulin was used as a loading control. **i** Western blot analysis of EXOC1 protein levels in HepG2-NTCP cells transfected with scrambled control (Ctrl) or *EXOC1*-targeting siRNAs. Cells were harvested at 2dpt/0dpi or following HBV infection at 9dpt/7dpi. β-tubulin was used as a loading control. **j-l** HBV infection readouts in HepG2-NTCP cells following siRNA-mediated *EXOC1* KD. Secreted HBeAg levels **(j)**, HBcAg-positive cells **(k)**, and extracellular HBV DNA levels **(l)** were quantified 7 dpi. Data in **b**, **d**, **f**, **g**, **j**, **k**, and L are presented as mean ± S.D. Statistical significance was assessed using two-tailed unpaired t-tests with Welch’s correction. \*\*\**P* < 0.001; \*\*\*\**P* < 0.0001.

### Identification and validation of *EXOC1* as an antiviral HBV host factor

*EXOC1* KO reproducibly increased HBcAg positivity across two independent HBV infections and exhibited the most pronounced enhancement of HBV infection markers among the validated candidates, making it the most compelling hit for subsequent investigation (Fig. 4a-d). *EXOC1* is a core component of the exocyst complex, a conserved regulator of intracellular vesicle trafficking^35^, providing an opportunity to explore whether host trafficking pathways contribute to HBV infection and replication. Western blot analysis confirmed reduced EXOC1 protein levels in *EXOC1*-targeted HepG2-NTCP cells, although *EXOC1* depletion was incomplete (Fig. 4e).

To further validate the antiviral role of *EXOC1* and assess whether it influences post-entry stages of the HBV life cycle, we examined *EXOC1* disruption in additional experimental systems. In HepG2-NTCP cells, transient delivery of recombinant Cas9-sgRNA ribonucleoprotein (RNP) complexes targeting *EXOC1* increased HBcAg positivity and secreted HBeAg levels following HBV infection relative to NT controls (Fig. 4f,g). Similarly, in HepDE19 cells^36^, which harbor an integrated HBV genome and therefore bypass viral entry, *EXOC1* targeting increased secreted HBeAg levels and showed a trend toward increased extracellular HBV DNA levels (Supplementary Fig. 2b,c). In both models, western blot analysis confirmed only partial depletion of EXOC1 protein (Fig. 4h; Supplementary Fig. 2d). Attempts to generate clonal *EXOC1* complete KO populations were unsuccessful.

Given prior reports of *EXOC1* being essential in other cell types^35,37–39^ and the inability to achieve a complete KO in hepatocyte cell lines, we reasoned that complete *EXOC1* loss may not be tolerated in proliferating cells and could generate heterogeneous cell populations that complicate the interpretation of HBV infection and replication phenotypes. We therefore employed an siRNA-mediated knockdown (KD) approach to achieve a more homogeneous reduction in EXOC1 protein expression. HepG2-NTCP cells were transfected with *EXOC1*-targeting or scrambled control siRNAs and infected with HBV 2 days post-transfection (dpt). EXOC1 protein levels were assessed by western blot both at the time of infection, 2 dpt, and 7 days post-infection (dpi), the experimental endpoint. Notably, siRNA-mediated knockdown achieved more robust EXOC1 depletion, which was maintained throughout HBV infection (Fig. 4i).

Consistent with the partial KO studies, *EXOC1* KD significantly increased all measured HBV readouts in HepG2-NTCP cells, including intracellular HBcAg, secreted HBeAg, and extracellular HBV DNA (Fig. 4j-l), supporting an antiviral role of *EXOC1* during HBV infection. To further demonstrate whether *EXOC1* also restricts HBV replication independently of *de novo* viral entry, we performed parallel KD experiments in HepDE19 cells. Efficient EXOC1 protein depletion was confirmed by western blot analysis at both early and late time points following siRNA transfection (Supplementary Fig. 2e). Similar to the phenotype observed in HepG2-NTCP cells, *EXOC1* KD in HepDE19 cells increased secreted HBeAg as well as extracellular HBV DNA levels relative to control siRNA-treated cells (Supplementary Fig. 2f,g).

Together, these findings establish *EXOC1* as a reproducible antiviral HBV host factor whose depletion enhances HBV infection- and replication-associated readouts across multiple experimental systems and independent perturbation strategies. The inability to achieve complete EXOC1 protein loss, combined with prior evidence that *EXOC1* is important for cellular fitness, further suggests that *EXOC1* may influence HBV infection through broader effects on host cellular pathways. Whether *EXOC1* directly restricts HBV replication or instead acts indirectly by altering host cellular programs remained unclear.

### *EXOC1* depletion induces hypoxia-associated and stress-response pathways during HBV infection

To determine whether *EXOC1* depletion alters broader host cellular programs in the context of HBV infection, we next performed transcriptomic profiling of *EXOC1*-depleted cells, in the presence and absence of HBV. HepG2-NTCP cells were transfected with *EXOC1*-targeting or NT control siRNAs and infected with HBV 2 dpt. Efficient EXOC1 depletion was confirmed by western blot analysis prior to infection and at the experimental endpoint (Fig. 4i). Cells were harvested at 7 dpi for differential gene expression analysis by RNA-sequencing.

Principal component analysis (PCA) revealed that *EXOC1* depletion was the primary driver of transcriptional variation across samples, whereas HBV infection contributed comparatively little to the overall transcriptomic differences observed between groups (Supplementary Fig. 3a). Control mock- and HBV-infected samples clustered closely together, as did *EXOC1*-depleted mock- and HBV-infected samples, indicating that *EXOC1* depletion induces broad transcriptional remodeling largely independent of HBV infection. Differential gene expression analysis comparing *EXOC1*-depleted and control HBV-infected cells identified numerous significantly upregulated and downregulated genes (Supplementary Fig. 3b-d), demonstrating extensive reprogramming of host cellular pathways following *EXOC1* depletion.

Because *EXOC1* functions as a core component of the exocyst complex^35^, we next examined whether depletion of *EXOC1* affected the expression of other exocyst family members. As expected, *EXOC1* expression was significantly reduced in *EXOC1*-depleted cells. Interestingly, *EXOC7* expression was also significantly decreased, likely not due to sequence similarity, whereas the remaining exocyst components showed little or no change (Fig. 5a), suggesting that *EXOC1* depletion may selectively influence the stability or regulation of specific exocyst complex subunits.

**Fig. 5.**
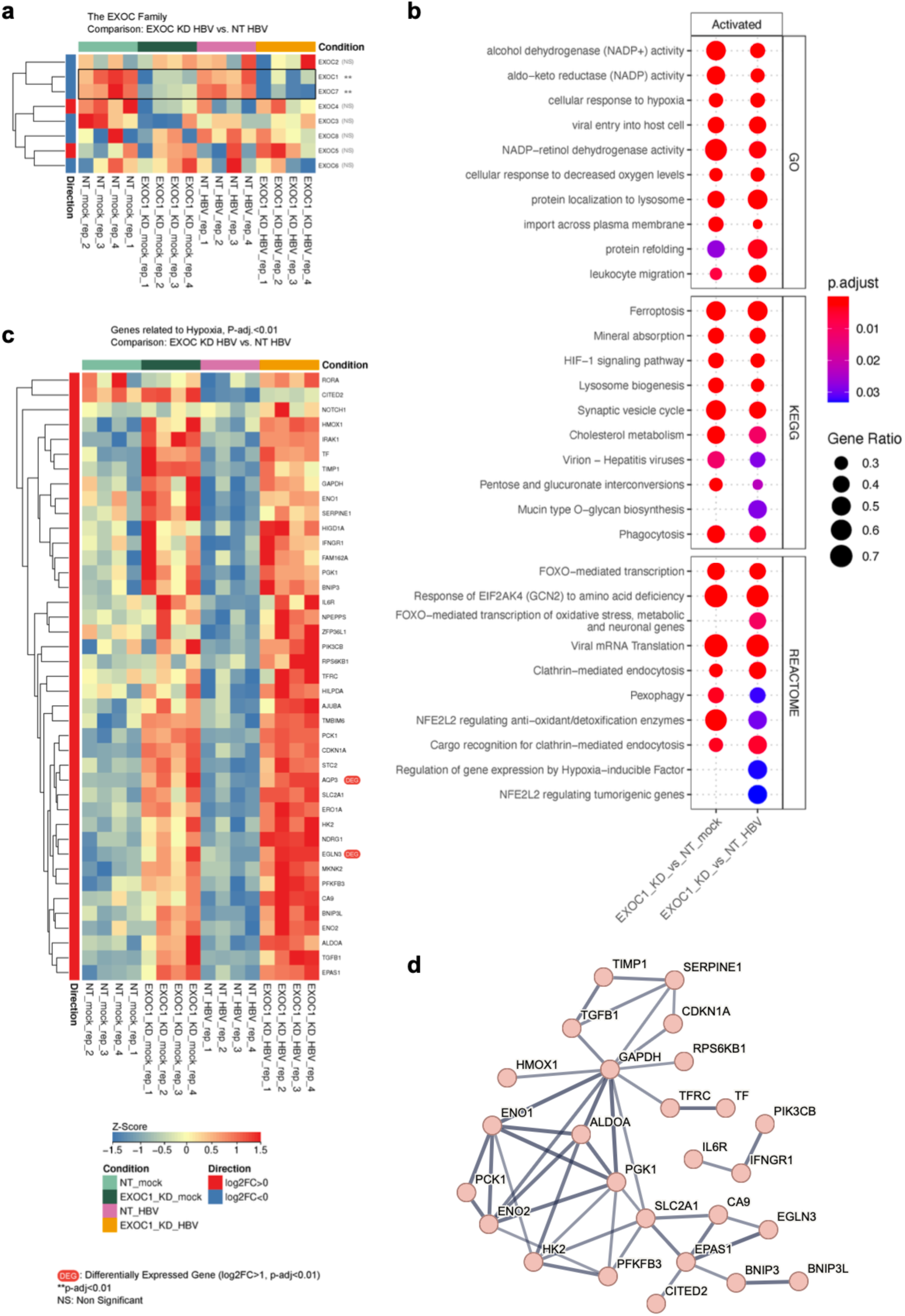
Transcriptomic profiling of EXOC1-depleted cells reveals enrichment of hypoxia-associated pathways during HBV infection. **a** Heatmap showing the expression of exocyst complex family members in control and *EXOC1* KD HepG2-NTCP cells in the presence and absence of HBV infection. *EXOC1* depletion resulted in a significant reduction of *EXOC1* expression and was accompanied by decreased *EXOC7* expression, whereas the remaining exocyst family members were largely unchanged. Statistical significance is indicated as \*\**P* < 0.01; NS, not significant. **b** Gene set enrichment analysis (GSEA) of genes differentially expressed following *EXOC1* depletion in mock-infected and HBV-infected cells. The top-enriched pathways (NES > 1.8) from Gene Ontology (GO), KEGG, and Reactome databases are shown. Dot size represents gene ratio, and color indicates adjusted *P*. **c** Heatmap of significantly upregulated genes (adjusted *P* < 0.01) associated with hypoxia-related pathways identified by GSEA in *EXOC1* KD HBV-infected cells relative to control HBV-infected cells. Gene expression values are shown as row-scaled Z-scores. **d** STRING network analysis^48^ of significantly induced hypoxia-associated genes shown in panel c. Only high-confidence interactions (interaction score ≥ 0.7) are displayed. Line thickness corresponds to interaction confidence. The network highlights interconnected genes involved in hypoxia signaling, metabolic adaptation, oxidative stress responses, and cellular survival pathways. The data represent four independent biological replicates per condition.

To further define the biological pathways altered by *EXOC1* depletion, we performed gene set enrichment analysis. This analysis identified significant enrichment of pathways associated with hypoxia, HIF-1 signaling, oxidative stress responses, ferroptosis, endocytosis, vesicle trafficking, lysosomal organization, and protein localization. Notably, hypoxia-associated pathways emerged among the most significantly enriched signatures across multiple databases, suggesting that *EXOC1* depletion reshapes the host cellular environment toward a hypoxic and metabolically stressed transcriptional state (Fig. 5b). In contrast, pathways suppressed following *EXOC1* depletion were overwhelmingly associated with cell-cycle progression, mitotic cell-cycle regulation, chromosome segregation, spindle organization, and DNA replication (Supplementary Fig. 3e). The coordinated downregulation of proliferation- and mitosis-associated pathways is consistent with previous reports implicating *EXOC1* in cellular homeostasis and proliferative capacity and may help explain the inability to generate complete *EXOC1* KO populations^35^.

Given the consistent enrichment of hypoxia-related pathways, we next focused on genes contributing to these signatures. Integration of significantly induced genes across hypoxia-associated gene sets identified 41 genes that were significantly upregulated following *EXOC1* depletion (adjusted *P* < 0.01) (Fig. 5c). These included multiple established hypoxia-responsive genes involved in oxygen sensing, metabolic adaptation, and cellular stress responses, including *EPAS1, EGLN3, CA9, SLC2A1, HK2, PFKFB3, ENO1, ENO2, PGK1, BNIP3, BNIP3L, HMOX1, SERPINE1*, and *CITED2*. STRING network analysis revealed extensive functional connectivity among these genes, forming a highly interconnected network centered on hypoxia signaling, glycolytic metabolism, oxidative stress adaptation, and cellular survival pathways (Fig. 5d).

Previous studies have linked hypoxia and HIF-dependent signaling to enhanced HBV transcription and replication^40,41^. The robust activation of hypoxia-responsive transcriptional programs following *EXOC1* depletion therefore provides a potential mechanistic explanation for the enhanced HBV phenotype in *EXOC1*-deficient cells. Given the established role of *EXOC1* in exocyst-mediated vesicle trafficking^35^, we speculate that disruption of EXOC1-dependent trafficking pathways may induce stress-associated transcriptional remodeling that converges on hypoxia-related signaling networks. Although the precise mechanistic relationship between *EXOC1* depletion, hypoxia-associated pathways, and HBV replication remains to be determined, these findings suggest that disruption of *EXOC1*-dependent trafficking pathways induces broad cellular remodeling characterized by activation of hypoxia-associated and stress-response programs, which may contribute to increased HBV permissiveness. Overall, transcriptomic profiling identified hypoxia-associated and stress-response pathways as prominent consequences of *EXOC1* depletion and highlighted these programs as potential contributors to the enhanced HBV replication phenotype.

### Human liver chimeric mice to assess HBV proviral factors *in vivo*

In the bulk KO studies, *EXOC1* was a strong antiviral factor, but because *EXOC1* depletion impairs cellular fitness and the human FNRG (huFNRG) model requires hepatocyte proliferation, *EXOC1* was not a tractable *in vivo* target (Supplementary Fig. 4a). We therefore focused our *in vivo* efforts on three proviral factors: *IRF2*, *WDR48*, and *ZCCHC14.* Using these genes, we established a human liver chimeric mouse platform to test whether their cell culture phenotypes extend to primary human hepatocytes *in vivo* (Fig. 6a, Supplementary Table 6), a setting that supports robust HBV replication and spread.

**Fig. 6.**
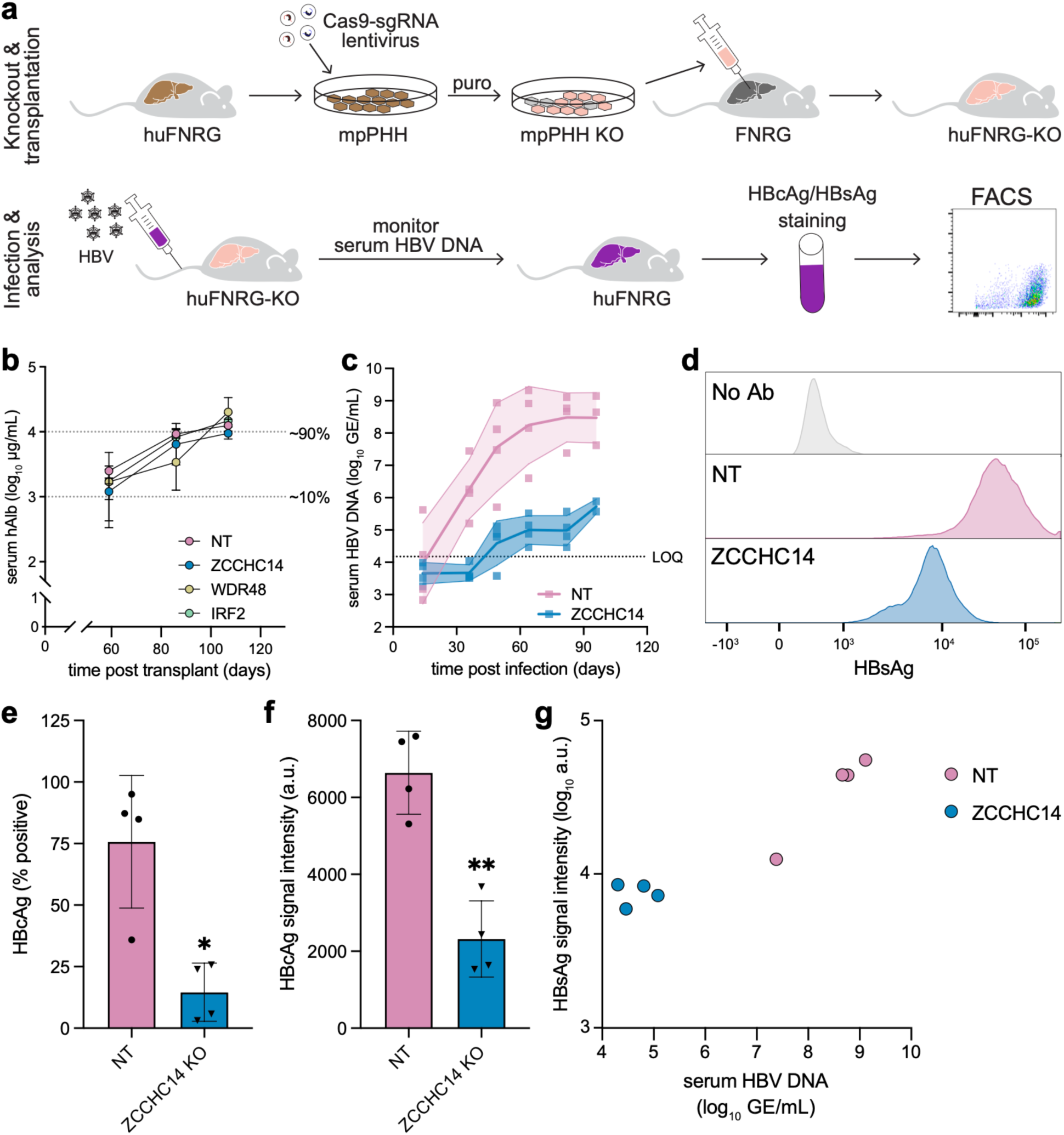
*In vivo* method for evaluating the effects of proviral host factors on engraftment and HBV infection. **a** Schematic of the generation of human liver chimeric mice with hepatocyte gene KOs followed by HBV infection and FACS-based analysis. **b** Serial serum hAlb levels of FNRG mice following transplantation of mpPHHs with NT or gene-targeting sgRNAs delivered by lentiviral transduction. Each symbol is the mean; the bars represent the range across mice in each group. Mice per group: n = 6 (NT, *IRF2* KO), n = 4 (*ZCCHC14* KO), n = 3 (*WDR48* KO). **c** Serial serum HBV DNA levels following humanization of FNRG mice are shown in (**a)** of NT mice and *ZCCHC14* KO mice. The dark line represents the mean for all mice per group; shading represents S.D.; each symbol represents an individual mouse and time point. Mice per group: n = 6 (NT), n = 4 (*ZCCHC14* KO). **d** Histograms of cell-associated HBsAg signal using FACS. Isolated mpPHHs from a representative isolation. Each histogram represents data from an individual mouse. No Ab, no primary antibody. **e** Percent of HBcAg-positive isolated mpPHHs measured by FACS. **f** HBcAg signal of HBcAg-positive isolated mpPHHs measured by FACS. **g** Serum HBV DNA (x-axis) is plotted against mean HBsAg signal intensity (y-axis). Each symbol represents an individual mouse; all mice from all isolations are included. For **d**-**g**, one million isolated mpPHHs from the HBV-infected, NT mouse, including in each isolation, were used for a no-primary-antibody control to establish the HBsAg and HBcAg-negative gates. Data in **e** and **f** are presented as mean ± S.D. Each symbol represents an individual mouse. Statistical significance was assessed using two-tailed unpaired t-tests with Welch’s correction. \**P* < 0.05; \*\**P* < 0.01.

We developed methods to generate KOs in mouse-passaged primary human hepatocytes (mpPHHs) *ex vivo*. KO mpPHHs were then transplanted into mice, and we monitored successful engraftment via serum human albumin (hAlb) levels, with ∼10 mg/mL corresponding to ∼90% humanization^42^. Although the kinetics of humanization varied depending on KO condition, by approximately 3 months post-transplantation, human-liver chimeric mice with *IRF2*, *WDR48*, or *ZCCHC14* KO hepatocytes reached humanization levels similar to the NT control mice, with hAlb levels corresponding to at least ∼90% humanization (Fig. 6b, Supplementary Fig. 4b, Supplementary Table 6).

Following successful human engraftment, human-liver chimeric mice were infected with HBV. HBV infection was monitored over time by measuring HBV DNA levels in the serum. Supporting previous studies of *ZCCHC14* as a HBV pro-viral factor in cell culture^18^, serum HBV DNA levels in the *ZCCHC14* human liver KO mice were approximately 3.5 logs lower than those of NT control mice (Fig. 6c). In contrast, we observed no effect of the *IRF2* KOs on serum HBV DNA levels (Supplementary Fig. 4c). *WDR48*’s effect was more modest, but on average the HBV DNA was reduced in the *WDR48* KO mice compared to NT control mice (day 64, 3 mice per group, mean NT = 4.8e8 GE/mL, mean *WDR48* KO = 7.8e6 GE/mL) (Supplementary Fig. 4d).

In addition to measuring serum HBV DNA levels, we isolated mpPHHs at peak viremia to evaluate cellular levels of HBsAg and HBcAg by FACS. The isolations were performed in four batches, with an NT and a *ZCCHC14* KO mouse included in each batch. We also confirmed that the isolation protocol yielded minimal mouse hepatocyte carryover, which could have overestimated the HBV-negative population (Supplementary Fig. 4e). At least half a million cells were analyzed per mouse and all were HBsAg-positive; the HBsAg signal intensity mirrored the observed serum HBV DNA levels (Fig. 6d). The percentage of HBcAg-positive cells, as well as the HBcAg signal intensity, was reduced in the *ZCCHC14* KO mice compared to NT control mice (Fig. 6e,f). We observed no differences in HBcAg positivity or levels between *IRF2* and *WDR48* KO mice and NT control mice (Supplementary Fig. 4f,g). HBV DNA levels showed the strongest correlation with the HBsAg signal intensity, R^2^ = 0.8028, *P* < 0.001 (Fig. 6g, Supplementary Fig. 4h,i).

In sum, three proviral targets were tested *in vivo,* and we observed no defects in engraftment in the human-liver chimeric mouse model. *ZCCHC14* KO showed a reproducible, strong reduction in serum HBV DNA, as well as reduced HBsAg and HBcAg levels, in isolated mpPHHs. *WDR48* exhibited a proviral role in the bulk KO experiments that was supported by a small reduction in serum HBV DNA levels *in vivo,* while *IRF2* did not show an effect in this model. Together, these studies validated *ZCCHC14* as a key proviral factor *in vivo* and established a human liver chimeric mouse platform for functional interrogation of HBV host factors.

## Discussion

Comprehensive genetic perturbation screens have become a powerful way to dissect virus–host interactions, illuminating host-factor networks across many viral families^15^. For HBV, however, the low efficiency of cell-culture infection and the scarcity of replication markers compatible with pooled selection have limited their use. Previous HBV screens have therefore relied on host-factor overexpression, HBV integration/replication systems, reporter viruses, or arrayed siRNA approaches^17–20^. Here, we performed, to our knowledge, the first pooled genome-wide CRISPR-Cas9 KO screen using *de novo* HBV infection and built a multi-step pipeline that advanced candidate host factors from orthogonal pooled screens to arrayed cell-line validation and *in vivo* testing in human liver chimeric mice. Three themes from this work merit further discussion.

First, the apparent HBV host-factor repertoire depends strongly on the experimental context in which it is measured. Overlap among HBV screens was modest, likely reflecting differences in perturbation strategies, cell types, viral systems, and readouts. This context dependence also persisted through validation. Of the 72 candidates carried forward, 38 genes (52.8%) were validated in at least one arrayed KO assay, yet the identities of validated genes differed depending on whether HBV was initiated by *de novo* infection or pgRNA transfection. For example, *AMBRA1* and *RABIF* were validated only in the infection-based arrayed assay and not after pgRNA transfection, a pattern consistent with roles at or before viral entry, although neither gene has an established entry-related function and both have instead been linked to autophagy or hepatocellular carcinoma^43,44^. Because these KOs did not alter cell number in either cell line, such infection-specific effects may also reflect cell-type differences between HepG2-NTCP and Huh7.5 cells, as previously observed for *CDKN2C*^17^. Rather than dismissing this divergence as technical variabiity, we interpret it as informative: perturbation modality, cell system, assay design, and entry dependence each shape which factors emerge, and triangulating across these contexts provides a more reliable route to robust HBV host factors than any single screen.

Second, this pipeline converged on a set of high-confidence candidates that included both known and previously unidentified HBV host factors. Among the proviral factors were known HBV dependency factor *ZCCHC14*^18^, along with *IRF2*, *WDR48*, and *FOXK1. EXOC1* emerged as the strongest antiviral factor. Confidence in the *EXOC1* phenotype was strengthened by the broader screen architecture: six of the eight core exocyst complex members were significant hits in the pooled screen, pointing to exocyst-mediated vesicle trafficking as one of the most prominent pathways in our dataset. This enrichment argues that the phenotype reflects a coherent biological process rather than an isolated gene-specific effect. Although exocyst components have been implicated in the replication of other RNA and DNA viruses^23–25^, their role in HBV infection had not yet been described.

*EXOC1* reduction enhanced HBV infection- and replication-associated readouts across *de novo* infection, HBV pgRNA transfection, and HepDE19 systems, suggesting that *EXOC1* influences post-entry stages of the HBV life cycle. Notably, this phenotype emerged despite only partial *EXOC1* depletion across multiple genome-editing strategies, and clonal *EXOC1* complete KO populations could not be recovered. Together with prior evidence that *EXOC1* is essential in other cell types^35,37–39^, these findings suggest that complete *EXOC1* loss may not be tolerated in proliferating hepatocyte lines and that even modest reductions in EXOC1 protein expression are sufficient to increase HBV permissiveness. More broadly, this highlights an important caveat of CRISPR KO-based host-factor discovery: efficient delivery of Cas9 and sgRNAs does not necessarily translate into complete protein depletion, particularly for genes linked to cellular fitness. Complementary approaches, such as CRISPR interference (CRISPRi)^15^ or tunable CRISPRi^45^, could enable the identification of essential host factors whose partial suppression is sufficient to impair HBV replication but whose complete loss is not tolerated, making them difficult to identify using conventional knockout screens.

How *EXOC1* restricts HBV remains to be established, but our data favor an indirect, host-centered mechanism rather than direct restriction of the viral replication machinery. Transcriptomic profiling of *EXOC1*-depleted cells revealed broad remodeling of host cellular programs, including reduced *EXOC7* expression, suppression of proliferation- and mitosis-associated pathways, and activation of hypoxia- and HIF-associated transcriptional signatures. The hypoxia signature is particularly notable because hypoxia and HIF-dependent pathways have previously been linked to enhanced HBV replication and viral transcription^40,41^. We therefore speculate that disruption of *EXOC1*-dependent trafficking shifts cells toward a hypoxia-like, metabolically stressed state that is more permissive to HBV. Future studies will be required to determine how exocyst function connects to hypoxia-associated signaling and whether this transcriptional state directly contributes to the enhanced HBV phenotype.

Third, extending validation into human liver chimeric mice helped prioritize which factors robustly affect HBV in primary human hepatocytes, a setting that cell lines do not fully recapitulate. By generating KOs in mpPHHs and engrafting them into FNRG mice^42,46^, we could test whether selected proviral candidates retained activity in a model that supports HBV infection and spread. The *in vivo* data sharpened our interpretation of the *in vitro* hits as *ZCCHC14* remained our strongest proviral target. *ZCCHC14* KO led to a marked reduction in serum HBV DNA and reduced viral antigen levels in isolated human hepatocytes, extending prior cell-culture studies of the ZCCHC14-TENT4A/B complex^18,47^ and supporting a robust proviral role for *ZCCHC14 in vivo*. In contrast, *WDR48* KO showed a more modest reduction in serum HBV DNA, consistent with its smaller effect in cell culture, whereas *IRF2* KO showed no detectable effect in this model.

These results raise an important question for future HBV host-factor validation: does the magnitude or reproducibility of an *in vitro* phenotype predict whether a candidate is worth pursuing *in vivo*? The strong concordance for *ZCCHC14*, the modest effect of *WDR48*, and the lack of detectable *in vivo* effect for *IRF2* are consistent with this possibility, but the number of genes tested here is too small to define a general principle. Systematic validation of additional hits will be needed to determine which features of *in vitro* phenotypes best predict *in vivo* relevance. *EXOC1* also illustrates a practical constraint of this strategy: because EXOC1 protein depletion appears to impair cellular fitness, it was not a tractable candidate for *in vivo* KO in a repopulating hepatocyte model, leading us to prioritize proviral factors for these studies. Encouragingly, cell-associated HBsAg measured by FACS correlated closely with serum HBV DNA, suggesting that small-scale pooled *in vivo* screens based on sorting and sgRNA sequencing may be feasible in future studies.

In summary, by combining orthogonal pooled and arrayed CRISPR-Cas9 KO screens with both *in vitro* and *in vivo* validation, we expand the catalog of HBV host factors and provide a framework and a resource for future host-directed target discovery in HBV. More broadly, our study highlights several lessons for genetic screening in HBV systems: candidate host factors are highly context dependent, editing efficiency does not always predict protein depletion or biological effect, and *in vivo* validation can substantially refine the prioritization of host factors identified in cell culture. Together, the insights gained and candidate host factors identified in this study advance our understanding of the HBV-host interaction landscape and will help guide future mechanistic investigations.

## Acknowledgments

We thank Dr. Margaret McDonald and other members of the Rice and Michailidis laboratories for their helpful discussions. We also thank the Rockefeller University‘s Comparative Bioscience Center and Flow Cytometry Resource Center. This work was supported by a sponsored research agreement from Vir Biotechnology Inc. (C.M.R., E.M.); NIH grants R21AI176944, R21AI176944-S1, R01AI181682, R56AI182395, DP1DA066168, and DP1DK139804 (to E.M.); R01AA027327 and P01HL160472 (to Y.P.J.); R01AI43295 and R01AI150275 (to C.M.R.); R01AI190067 (to C.M.R., Y.P.J., and E.M.); Stavros Niarchos Foundation (SNF) as part of its grant to the SNF Institute for Global Infectious Disease Research at the Rockefeller University (to C.M.R.). C.A.F. is currently supported by NIH/NIAID MOSAIC award K99AI190315 and was the Berger Foundation Fellow of the Damon Runyon Cancer Research Foundation (DRG-2440-21). J.B. was supported by FIOCRUZ and CNPq 312671-2022-2, 205096-2018-2. A.L.P.M. was supported by FIOCRUZ and CAPES 88881.337176/2019-01. X.H. is an Open Philanthropy Awardee of the Life Science Research Foundation and was supported by a C.H.Li Memorial Scholar Fund Award at The Rockefeller University.

## Author contributions

C.A.F., G.D., A.P., F.A.L., H.W.V., S.H., W.M.S., Y.P.J., C.M.R., and E.M. conceptualized the study. C.A.F., G.D., A.A., E.M., M.K.S., A.H.D.C., C.K.B., D.L., M.B., S.L., G.C., G.L., A.L.P.M., J.B., K.K., L.F., C.Z., Y.Z., C.Q., L.L.S., X.H., A.R., and Y.P.J. performed the experiments. C.A.F., G.D., A.A., E.M., M.B., L.B.S., A.T., and J.I. performed the analysis. Y.Y., A.P., F.A.L., H.W.V., S.H., W.M.S, Y.P.J., C.M.R., and E.M. supervised the research. C.A.F., G.D., A.A., M.B., K.K., and Y.P.J. completed the visualization of the results. C.A.F. and G.D. wrote the original draft of the manuscript. All authors reviewed and edited the draft.

## Materials and methods

### Cells

HepG2-NTCP cells^49^ were cultured in Dulbecco’s Modified Eagle Medium, DMEM (Sigma-Aldrich #D6429) supplemented with either 3% or 10% fetal bovine serum, FBS (Cytiva #SH3030396 or HyClone Laboratories) and 0.1 mM non-essential amino acids, NEAA (Thermo Fisher Scientific #11140050) and in some cases with 1% Anti-Anti (Thermo Fisher Scientific #15240062) at 37°C and 5% CO_2_.

Huh7.5-Cas9 cells were cultured in DMEM (Thermo Fisher Scientific #11995065), supplemented with 0.1 mM NEAA (Thermo Fisher Scientific #11140076) and 10% FBS (HyClone Laboratories) at 37°C and 5% CO_2_.

HEK293-pLentiX cells were cultured in DMEM (Sigma-Aldrich #D6429) supplemented with 10% FBS (Cytiva #SH3030396) and 0.1 mM NEAA (Thermo Fisher Scientific #11140050) at 37°C and 5% CO_2_.

HepDE19 cells harboring a tetracycline-regulated (Tet-off) HBV expression system were cultured in DMEM supplemented with either 3% or 10% FBS and 0.1 mM NEAA. Where indicated, tetracycline was included (+Tet) or omitted (−Tet) to suppress or induce HBV expression, respectively.

All cell lines tested negative for mycoplasma.

### Animal studies and primary hepatocytes

#### Mice

*Fah^-/-^ NOD Rag1^-/-^ Il2rg^null^* (FNRG) mice were generated by backcrossing *Fah^-/-^* liver injury mice^50^ to *NOD Rag1^-/-^ Il2rg^null^*(NRG, Jackson Laboratories) animals for 13 generations^51^. To create human liver chimeric FNRG mice, animals were preconditioned with retrorsine and withdrawn from nitisinone (Yecuris) prior to transplantation with 5 × 10^5^ primary human hepatocytes from donor HUM4188 (Lonza)^42^. Animals were cycled on nitisinone for 10-12 weeks, after which human albumin (hAlb) was quantified in mouse serum using a species-specific ELISA (Bethyl Labs). Animals with hAlb levels >9 mg/mL were used for infection experiments. All procedures were approved by the Rockefeller University IACUC under protocols 18063, 21056, and 24022.

#### Mouse-passaged primary hepatocytes (mpPHH)

The primary hepatocytes used in this study were derived from human donors and expanded in liver-humanized mice. Mouse-passaged primary human hepatocytes (mpPHH) were isolated from highly humanized FNRG (huFNRG) mice as previously described^42^. Unless otherwise indicated, experiments were performed using mpPHH derived from a single donor (#HUM4188) to minimize variability and enable controlled comparisons between conditions.

In brief, humanized FNRG mice were anesthetized with ketamine/xylazine, and a 24-gauge angiocath was inserted into the inferior vena cava. Then, the portal vein was cut, and the mouse liver was perfused sequentially with 1× PBS^−/−^ supplemented with heparin, Hanks buffered saline solution (HBSS) supplemented with 5 mM ethylenediaminetetraacetic acid (EDTA) and 50 mM 2-[4-(2-hydroxyethyl)piperazin-1-yl] ethanesulfonic acid (Hepes), and finally, HBSS supplemented with 0.05% Type IV collagenase (Sigma-Aldrich) and 1 U/mL DNase (Thermo Fisher). After digestion, the liver was disrupted over a 70-μm cell strainer, and the cell suspension (50 mL) was centrifuged at 50 × g for 5 min at 4°C using an Allegra X-14R Centrifuge (Beckman Coulter). The supernatant was gently aspirated, and the cells were washed once with 1× PBS^−/−^. The cell pellet was resuspended in 10 mL 1× PBS^−/−^ and gently mixed with an equal volume of Percoll working solution (GE Healthcare #17-0891-01). Percoll working solution consisted of 5.4% 10× PBS^−/−^, 48.6% Percoll, and 46% William’s E medium (Life Technologies #12551-032). The cell suspension was centrifuged at 100 × g for 5 min at 4°C, and the pellet was washed once with 1× PBS^−/−^. After centrifugation at 50 × g, the cells were resuspended in 1× PBS^−/−^ and left on ice for 45 min. A second Percoll purification was then performed to further purify the cell suspension. The final pellet was resuspended in W10 plating medium. W10 medium is William’s E medium supplemented with 10% FBS, 1% penicillin/streptomycin (Life Technologies #15140–12), 1% 200 mM L-glutamine (Life Technologies #25030-081), 0.1% Gentamicin reagent solution (50 mg/mL) (Life Technologies #15750-060), and 0.1% Corning ITS premix (Corning #354350). Viable cells were counted using trypan blue. Cell viability was typically above 95%, and minimal cell clumping was observed. When cell clumps were present, the suspension was filtered through a 40 μm cell strainer.

For mpPHH culture and maintenance, cells were kept in HCM medium (Lonza #CC3198) supplemented with 2% DMSO, 1% gentamicin (Gibco #15750–060), and 0.5% ciprofloxacin (Sigma-Aldrich #17850-5G-F). Media was typically replaced every 2-3 days. Distinct mpPHH mouse numbers throughout the study refer to independent hepatocyte preparations isolated from separately humanized FNRG mice engrafted with hepatocytes originating from the same human donor unless otherwise indicated.

### Viruses

All CRISPR-Cas9 pooled knockout (KO) screens used an HBV genotype D stock from HepAD38 cells that was either produced using the method below or purchased from Imquest Biosciences. HepAD38 cells were maintained in HepAD38 culture medium with tetracycline at 37°C and 5% CO_2_ for two weeks. Upon reaching confluency, cells were switched to media without tetracycline (−Tet) to induce HBV replication. Starting at day 14 post-tetracycline removal, supernatant was collected, centrifuged, and filtered to remove cell debris. Virus-containing supernatant was polyethylene glycol (PEG)-precipitated with a final PEG concentration of 6.5%. Concentrated virus stocks were aliquoted and stored at −80°C until use.

HBV used for validation and subsequent experiments was produced from HepDE19 cells, as described previously^36^. Prior to confluency, cells were maintained in DMEM supplemented with 10% FBS, 0.1 mM NEAA, and 1 μg/mL tetracycline. Upon reaching confluency, cells were switched to DMEM containing 3% FBS and 0.1 mM NEAA without tetracycline (−Tet) to induce HBV replication. Culture supernatants were collected every 2-3 days for up to 4 weeks, and virus-containing media were concentrated using Centricon Plus-70 centrifugal filter devices (Cat# UFC710008; MilliporeSigma). Concentrated viral stocks were aliquoted and stored at −80°C until use.

The virus used for *in vivo* experiments was generated as follows. The 1.31× HBV-A_A plasmid (kindly provided by Drs Amir Shlomai and Yossi Shaul, Weizman Institute, Israel)^52^ was digested with PciI (New England Biolabs (NEB) #R0655) and BglI (NEB #R0143), gel purified and ligated to create recombinant (r)cccDNA. After gel purification, 1 µg of rcccDNA was injected into the liver of huFNRG mice with human albumin serum levels around 10 mg/mL. Upon reaching plateau viremia, serum from these mice was used to inoculate naive huFNRG mice.

### CRISPR-Cas9 pooled knockout screens

The Human Brunello CRISPR knockout pooled library was a gift from David Root and John Doench (Addgene #73179-LV; http://n2t.net/addgene:73179-LV; RRID:Addgene_73179)^21^. The provided lentiviral library encoding both SpCas9 and unique sgRNAs for 19,114 genes was introduced into HepG2-NTCP cells via transduction at low multiplicity of infection, followed by selection. Selected cells were then expanded to reach sufficient coverage of the library before seeding to perform the screens described below.

#### HBsAg full-genome screen (HBs-FG)

Approximately 9 × 10^6^ cells were seeded per T175 flask. Twenty flasks were used for infected samples and 12 for uninfected controls. For infected samples, cells were infected with HBV genotype D (see stock details in Section “Viruses”) at 1,000 genome equivalents per mL (GE/mL) in the presence of 4% PEG overnight and 2.5% dimethyl sulfoxide (DMSO). All flasks were harvested 8 days post-infection (dpi). One set of samples included infected and mock treated cells without sorting. See below for staining, sorting, and DNA extraction method details.

#### HBcAg full-genome screen (HBc-FG)

Approximately 9 × 10^6^ cells were seeded per T175 flask. Fifty T175 flasks were infected with HBV genotype D at 1,000 GE/mL three times on three consecutive days in the presence of 4% PEG overnight and 2.5% DMSO. All flasks were harvested 8 days following the first infection.

#### HBcAg targeted screen (HBc-T)

A lentiviral library comprised of genes selected from the HBs-FG screen results (top 960 genes ranked by p-value) was generated. The targeted sgRNA library contained 3840 sgRNAs, approximately 4 sgRNAs per gene and was generated and obtained through the Genetic Perturbation Platform at the Broad Institute of Massachusetts Institute of Technology (MIT) and Harvard, using production methods described previously^21^. Forty million cells were seeded in a triple-layer T175 flask. As in HBc-FG, cells were infected with HBV genotype D at 1,000 GE/mL three times on three consecutive days. Cells were harvested 11 days post the first infection.

#### Screen staining and sample processing

For all three screens, cells were fixed in 4% paraformaldehyde (PFA). For HBs-FG, infected cells were stained overnight at 4°C for HBsAg using an in-house monoclonal primary antibody (HBC24 mAb, 1 µg/mL with 0.5% casein) and then with a secondary antibody for detection (mAb, anti-human 647). For HBc-FG, infected cells were stained overnight at 4°C for HBcAg using primary antibody (Austral HBc mAb, diluted 1:1,000 with 0.5% casein) and then with secondary antibody for detection (mAb, anti-rabbit 488). For HBc-T, cells were stained under the same conditions as the HBc-FG except with the HBc Ab (Cell Marque HBc mAb, diluted 1:1,000 with 0.5% casein). Stained samples were then sorted (Aria Fusion or Sony MA900) by HBsAg or HBcAg signal. Genomic DNA was extracted from all samples with the Qiagen FFPE kit (Qiagen, catalog no. 56404), according to the manufacturer’s instructions except for the following changes: 0.3 M NaCl was added to the lysis buffer, and incubation at 56°C was extended to 4 h. Sequencing libraries and next-generation sequencing were performed as previously described^21^.

### Analysis of screens and previously published screen data

To assess the relative abundance of sgRNAs, MAGeCK RRA was used^53^. The comparisons used to determine the enrichment or depletion of genes based on sgRNA counts are outlined below for each screen.

The comparisons for HBs-FG were as follows: (i) infected unsorted vs mock unsorted, (ii) top 10% of cells by HBsAg signal vs infected unsorted, and (iii) bottom 10% of cells by HBsAg signal vs infected unsorted. Selected genes from the HBs-FG screen were those with a *P* < 1e-03 (n = 236 genes) and no fold-change cut-off. For HBc-FG and HBc-T: (i) HBcAg negative versus infected unsorted, (ii) HBcAg positive versus infected unsorted, and (iii) HBcAg negative versus HBcAg positive.

Selected genes from the HBs-FG screen and the Brunello library gene list were uploaded to STRING-db^48^. Of the 236 genes, 229 were properly mapped to those contained within STRING-db. Using the Brunello library gene list as the background gene list, gene set enrichment analysis was performed using the following settings: no merging of gene sets by similarity, databases (GO-Process, GO-Function, KEGG, REACTOME), and remaining default parameters. Enriched gene sets listed in Supplementary Table 2 are those with FDR < 0.05, signal > 0.01, strength > 0.01, and 2 or more genes from the HBs-FG screen hits. In addition, a full STRING network was generated displaying only high-confidence interactions (score > 0.9), with edge widths proportional to confidence scores. Only those of the 229 genes with at least one functional interaction were displayed in the network.

For the published screen dataset comparison, the complete screen datasets were downloaded from the respective references. For the full-genome pooled, CRISPR-Cas9 KO screen with integrated HBV (Int-Cas9-pool)^18^, the gene list was filtered using the same min.RSA threshold of −4.96 as in the original study but no fold-change threshold (204 genes). For the full-genome arrayed, siRNA KD screen with HBV infection (Inf-siRNA-array)^20^, the same activity threshold for an active siRNA from the original study was used. The gene list was then filtered to include genes for which 3 of 3 siRNAs were classified as active, or 2 of 3 siRNAs were active and the fold-change directionality was consistent (1,003 genes). R (v4.5.0) software packages ggVennDiagram (v1.5.7) and GeneOverlap (v1.46.0) were used to visualize the gene overlap and perform statistical analyses.

Selected genes for arrayed validation experiments were those with a *P* < 1e-03 in at least two of three pooled screens and not related to p53 and heparan sulfate proteoglycans. Final selected gene count was 72 (see Supplementary Tables 4 and 5 for gene symbols).

### Arrayed CRISPR-Cas9 validation screen in HepG2-NTCP cells with HBV infection

The 72 selected genes identified from the screens were validated using an arrayed CRISPR-Cas9 KO approach in HepG2-NTCP cells. Cells were seeded in 96-well plates at 5,000 cells per well and edited with Cas9-sgRNA RNP complexes composed of recombinant Cas9 nuclease (Integrated DNA Technologies (IDT) #CAS12207) and synthetic sgRNAs (IDT). RNP complexes were assembled in Opti-MEM Reduced-Serum Media (Opti-MEM, Thermo Fisher Scientific #51985034) and delivered using DharmaFECT Duo transfection reagent (Horizon Discovery #T-2010-03) according to the manufacturer’s instructions. Final concentrations of Cas9 and sgRNA were 40 nM and 50 nM, respectively. NT and lethal sgRNA controls were included on every plate to monitor editing specificity and transfection efficiency, respectively. Following transfection, plates were centrifuged at 1,000 × g for 60 min at 37°C to enhance RNP delivery.

To support HBV infection, edited cells were cultured in DMEM supplemented with 3% FBS and 2% DMSO (ATCC #4-X) for 24 h prior to infection. Cells were infected with HBV in the presence of 4% PEG8000 (Sigma-Aldrich #81268) and 2% DMSO. Viral inoculum was added to the cells, followed by spinoculation at 1,000 × g for 1 h at 37°C. Cells were subsequently incubated with the inoculum for 24 h, then washed with 1× DPBS and maintained in medium containing 3% FBS and 2% DMSO. Media were replenished at 3 dpi, and HBV replication markers were assessed at 7 dpi, see details below.

### Arrayed CRISPR-Cas9 validation screen in Huh7.5-Cas9 cells with pgRNA transfection

Preparation of 5’ capped and 3’ polyadenylated pgRNAs was generated as previously described^32^. In brief, 10 µg of HBV genotype A plasmid DNA was linearized with NotI-HF (NEB #R3189S) and then purified with the MinElute PCR Purification kit (Qiagen #28004) according to the manufacturer’s protocol. Transcription was performed using 2 µg of linearized and purified plasmid as input and the RiboMax T7 kit (Promega #P1320) according to the manufacturer’s protocol. Reactions were incubated for 30 min at 37°C, followed by an additional 15 min at 37°C after adding 2 µL of Turbo DNase I (Thermo Fisher Scientific #AM2238). DNase-treated, transcription products were then purified using the RNeasy mini kit (Qiagen #74014) according to the manufacturer’s instructions, including the on-column DNase step (Qiagen #79254). Capping and polyadenylation were performed with 40-60 µg of purified RNA as input using the Cap 1 and polyadenylation reagents included in the T7 mScript Standard mRNA Production System kit (Cellscript #C-MSC116) according to the manufacturer’s instructions, with 30-min incubations at 37°C for each step. Capped and polyadenylated RNA was then purified with the RNeasy kit without the on-column DNase treatment.

In 96-well format, sgRNAs (IDT) were delivered to Huh7.5-Cas9 cells via reverse transfection. For each well, 0.05 µL of Dharmafect-4 (Horizon Discovery #T-2004) and 0.25 µL of a 250 nM sgRNA stock were diluted in Opti-MEM (Thermo Fisher Scientific #51985034) to a final volume of 20 µL. Dharmafect-4 and sgRNA mixtures were then incubated for 20 min at room temperature. During the incubation, Huh7.5-Cas9 cells were prepared to a concentration of 50,000 cells/mL (i.e., 4,000 cells were seeded per well) in Huh7.5-Cas9 growth media. For each well, 20 µL of sgRNA transfection mix was combined with 80 µL of Huh7.5-Cas9 cells, then centrifuged at ambient temperature for 5 min at 500 × g. The cells were then kept at 37°C and 5% CO_2_ until HBV pgRNA transfection.

Three days post sgRNA reverse transfection, cells were transfected with HBV pgRNA transfection using methods similar to those described previously^32^. Prior to transfection, the medium was replaced with DMEM supplemented with 0.1 mM NEAA and 1.5% FBS. For each well, 16.8 ng of HBV pgRNA was mixed with 0.084 µL of Lipofectamine 2000 (Thermo Fisher Scientific #11668-019) in a total volume of 8.5 µL Opti-MEM and incubated for 20 min at room temperature. This mixture was added to cells, followed by centrifugation at 37°C for 30 min at 1,000 × g and then returned to 37°C and 5% CO_2_. Six hours following transfection, the medium was removed and replaced with DMEM supplemented with 0.1 mM NEAA, 3% FBS, and 2% DMSO. At two and four days post pgRNA transfection, the supernatant was collected and replaced with fresh reduced serum and DMSO-containing media as defined above. The collected supernatant was stored at - 20°C until HBsAg detection assays were performed (see below for details). Four days post HBV pgRNA transfection, nuclei were stained with 1 µg/mL Hoechst 33342 (Invitrogen #H3570). Images were collected on the BioTek Cytation 7 Cell Imaging Multimode Reader (Agilent) and analysis performed by the Gen5 software (Agilent).

### Cellular HBcAg staining

Cells were fixed in 4% PFA for 20 min at room temperature, washed with 1× DPBS, and permeabilized with 0.1% Triton-X100 for 10 min. After washing with PBS, cells were incubated for 1 h at room temperature with 5% goat serum (Jackson ImmunoResearch) in PBS. Cells were incubated overnight at 4°C in a 1:1,000 dilution of rabbit polyclonal anti-HBV core antibody (Cell Marque #216A-16-ASR) in PBS with 5% goat serum (Sigma Aldrich). Cells were washed with PBS with 0.1% Tween-20 (Promega), then incubated for 1 h at room temperature with goat anti-rabbit AlexaFluor 594 (Life Technologies) at a dilution of 1:1,000. Cells were counterstained with 1:1,000 Hoescht (Thermo Fisher Scientific). Cells were washed twice with PBS-T and once with PBS before imaging using the BioTek Cytation 7 Cell Imaging Multimode Reader (Agilent) and analysis performed by the Gen5 software (Agilent).

### HBeAg and HBsAg chemilluminescence assays (CLIA)

Supernatant levels of HBeAg and HBsAg were quantified using the HBeAg and HBsAg chemiluminescence immunoassays (CLIA). Kits were purchased from DiaSino [#DS187702 (HBeAg) and #DS187703 (HBsAg)] or Ig Biotechnology [#CL18004 (HBeAg)]. Assays were performed according to the manufacturer’s protocol.

### Integrated analysis following arrayed validation

Using the arrayed validation results, we generated a list of genes for further assessment. This list of genes included those that were validated in at least one of the arrayed screening readouts with the following parameters, where thresholds were based on the values shown in Figs. 2b, d, 3b: HepG2-NTCP HBeAg (<0.75 or >1.5); HepG2-NTCP HBc-positivity (<0.75 or >1.25); Huh7.5-Cas9-pgRNA (<0.5 or >2). If a gene was significant in more than one arrayed screen, then the directionality of the effect had to be the same for the gene to be included. The list of genes is included in Supplementary Table 5.

In addition to the arrayed assays, the screen-identified genes (72 genes) were assessed using a literature-informed analysis described in brief. Genes hypothesized to be involved in entry, vesicle trafficking, mitochondria, and translation factors based on gene set categorization and gene function were filtered, resulting in a list of 28 genes. These 28 genes were then evaluated and scored on a series of metrics: known gene function, hypothesized HBV mechanism, gene essentiality (i.e., whether a KO would be toxic), and whether the gene is targeted by known drugs.

Combining the arrayed validation and 28 genes from the above analysis resulted in 53 genes (38 validated, 15 literature-informed, 13 overlapping) that were filtered based on an integrated analysis incorporating the validation results and metrics defined above to select 13 for bulk KO experiments. Graphical representation of this analysis workflow is shown in Supplementary Fig. 2a.

### LentiCRISPRv2 transduction

Lentiviral particles were produced in HEK293-pLentiX cells cultured in T175 flasks. Prior to transfection, culture medium was replaced with DMEM supplemented with 10% FBS and 0.1 mM NEAA. For each T175 flask, sgRNA or Cas9 transfer plasmids were co-transfected together with the second-generation packaging plasmid psPAX2 (5.5 μg; Addgene #12260) and envelope plasmid pMD2.G (2 μg; Addgene #12259) using FuGENE 4K transfection reagent (45 μL; Promega #E5911) in Opti-MEM (682 μL). Transfection mixtures were incubated at room temperature for 30 min before addition to the cells.

Viral supernatants were harvested at 48 and 72 h post-transfection, clarified by centrifugation, filtered through 0.45 μm filters, aliquoted, and stored at −80°C until use.

For *in vivo* experiments, mpPHH were transduced with lentiviral expression vectors. Plated mpPHHs were transduced in hepatocyte culture medium (HCM) supplemented with 2% DMSO and 4 μg/mL polybrene (Sigma-Aldrich #TR-1003-G) by spinoculation at 1,000 × g for 1 h at 37°C. For the NT control and *in vivo* tested genes except *ZCCHC14*, cells were co-transduced with a 1:1 mixture of two lentiviral vectors expressing two independently designed sgRNAs. For *ZCCHC14* targeting, a single designed sgRNA and one lentiviral stock were used to transduce mpPHHs. Following transduction, cells were maintained in HCM supplemented with 2% DMSO, with media changes every 2-3 days. Cells were harvested 14 days post-transduction to allow sufficient time for Cas9 and sgRNA expression, genome editing, and turnover of pre-existing target protein.

### Recombinant Cas3 and synthetic EXOC1 sgRNA transfection

Cells were transfected with recombinant Cas9 protein (IDT #CAS12207) complexed with synthetic sgRNAs. Synthetic sgRNAs and recombinant Cas9 were thawed on ice and combined in Opti-MEM at 37°C for 5 min for RNP formation. DharmaFECT Duo transfection reagent was added to the culture medium. Then, both suspensions were mixed at a 1:1 ratio and, incubated at room temperature for 20 min, and added directly to each well (for a 96-well plate: 50 µL/well). Final concentrations were 1% for DharmaFECT Duo, 60 nM for Cas9, and 100 nM for sgRNA. Sequentially, cells were centrifuged at 1,000 × g for 1 h at 37°C (spinoculation) to enhance uptake. Following transfection, cells were maintained in culture media supplemented with 2% DMSO, with medium changes every 2-3 days. NT sgRNAs were obtained from Horizon Discovery (Dharmacon Edit-R platform; #U-007501-01-20, U-006000-01-20, and U-002005-20). The exact guide RNA sequences are proprietary and were not disclosed by the manufacturer.

### HBV infection in bulk KO and siRNA transfected cells

HepG2-NTCP cells were seeded at a density of 25,000 cells per well in a 96-well collagen-coated plate in DMEM supplemented with 10% FBS and 0.1 mM NEAA. The following day, the media was changed to DMEM supplemented with 3% FBS, 0.1 mM NEAA, and 2% DMSO to reduce cell proliferation and enhance susceptibility to HBV infection. The next day, cells were infected with HBV derived from HepDE19 cells or virus-free control inoculum in the presence of 4% PEG8000 and 2% DMSO. Viral inoculum was added to the cells, followed by spinoculation at 1,000 × g for 1 h at 37°C. Cells were subsequently incubated with the inoculum for 24 h. The next day, the inoculum was removed, the cells were washed three times with 1× DPBS, and fresh DMEM supplemented with 3% FBS, 0.1 mM NEAA, and 2% DMSO was added^49^. Media were replenished at 3 dpi. At 7 dpi, culture supernatants were collected, and cells were fixed with 4% PFA for downstream analyses.

### Western blot analysis

Cells were washed with PBS, scraped, pelleted, and lysed with RIPA Lysis and Extraction Buffer (Thermo Fisher Scientific #89901), followed by separation by SDS-PAGE and transfer to nitrocellulose membrane. Membranes were incubated overnight in primary antibodies [HRP-conjugated Beta Tubulin Monoclonal antibody (Proteintech #HRP-66240), EXOC1 Polyclonal Antibody (Thermo Fisher Scientific #PA5-58071)] diluted in 0.1% milk/TBS-T. TBS-T was used to wash the membranes before incubation with secondary HRP-conjugated antibodies (Thermo Fisher Scientific #31460) of the corresponding species. Pictures were taken with an Azure imager.

### Measurement of HBV DNA

To quantify extracellular HBV DNA in the *in vitro* experiments, DNA was extracted from 200 μL of supernatant using the QIAamp DNA Blood Mini Kit (Qiagen #51106), and extracellular DNA was eluted in 50 µL of elution buffer. To quantify HBV DNA in the serum of huFNRG mice, 25 µL of serum was diluted in 175 µL of 1× DPBS, DNA was extracted using the QIAamp DNA Blood Mini Kit (Qiagen #51106), and serum DNA was eluted in 60 µL elution buffer. Finally, HBV DNA levels were measured by qPCR using a TaqMan-based assay, as previously described^32^.

### Gene knockdown using siRNAs

siRNA-mediated knockdown of *EXOC1* was performed using a pool of four siRNAs targeting *EXOC1* (siGENOME SMARTpool siRNA; Horizon Discovery #M-013312-01-0005). A siGENOME Non-Targeting siRNA Control Pool #2 (Horizon Discovery #D-001206-14-05) was used as a negative control. siRNA transfections were conducted using DharmaFECT-4 transfection reagent at a final concentration of 0.4% with siRNAs used at final concentrations of 50 nM, according to the manufacturer’s instructions.

#### RNA sequencing and analysis

HepG2-NTCP cells were transfected with NT or *EXOC1* siRNAs then infected with HBV as described above. Cells were trypsinized, pelleted, and resuspended in DNA/RNA Shield buffer (Zymo Research #R1100) according to the manufacturer’s instructions, and submitted to Plasmidsaurus for RNA-seq analysis.

Single-end reads were generated using the Illumina NovaSeq X Plus sequencing platform. Read quality control was assessed before and after trimming using FastQC v0.11.9. Adapter trimming and quality filtering were performed using fastp v1.3.1^54^, enforcing a minimum quality threshold of Q20 and a minimum read length of 36 bp. Trimmed reads were quantified using Salmon v1.11.4^55^ via selective alignment against a partial selective alignment index. This index comprised the human transcriptome (GENCODE v49) alongside the human primary assembly genome (GRCh38) and the HBV RefSeq (NC_003977.2) acting as decoys. Transcript-level quantifications were imported and aggregated to the gene level using the tximport library^56^ in R v4.3.2. Low-abundance genes were pre-filtered by retaining only those with a minimum of 10 counts in at least all 4 samples. Differential expression analysis was performed using DESeq2^57^. To control for variance and improve visualization, log_2_ fold-change (log_2_FC) values were shrunk using the Adaptive Shrinkage (ashr) method. Variance-stabilizing transformation (vst) was applied to the counts for downstream exploratory analyses, including principal component analysis and heatmap visualizations. Statistical significance was defined using a Benjamini-Hochberg adjusted P-value (FDR) threshold of less than 0.01 and an absolute log_2_FC greater than 1. Functional enrichment analyses were performed utilizing the clusterProfiler library^58^ incorporating annotations from the org.Hs.eg.db, ReactomePA^59^, and DOSE packages^60^. The input of genes for gene set enrichment analysis considered FDR values and not the additional log_2_FC threshold. All data visualizations were generated using ggplot2.

### Generation of mpPHH KOs for humanization followed by HBV infection

For *in vivo* experiments, mpPHH were transduced with lentiviral expression vectors. Plated mpPHHs were transduced in hepatocyte culture medium (HCM) supplemented with 2% DMSO and 4 μg/mL polybrene (Sigma-Aldrich #TR-1003-G) by spinoculation at 1,000 × g for 1 h at 37°C. For the NT control and *in vivo* tested genes except *ZCCHC14*, cells were co-transduced with a 1:1 mixture of two lentiviral vectors expressing two independently designed sgRNAs. For *ZCCHC14* targeting, a single designed sgRNA and one lentiviral stock was used to transduce mpPHHs. Following transduction, cells were maintained in HCM supplemented with 2% DMSO, with media changes every 2-3 days. Cells were harvested 14 days post-transduction to allow sufficient time for Cas9 and sgRNA expression, genome editing, and turnover of pre-existing target protein and engrafted using methods described above in section “Animal studies and primary hepatocytes.*”* Then, the huFNRG mice carrying KO or NT grafts were inoculated retroorbitally with serum from viremic mice containing 1 × 10^6^ genome equivalents HBV-A_A per mouse.

### Serum human albumin measurement

Serum was collected from the tail vein or retro-orbital plexus of humanized FNRG mice. Levels of hAlb were quantified using a sandwich ELISA assay (Bethyl Laboratories) to determine the extent of human hepatocyte repopulation, as serum hAlb levels correlate with liver humanization in FNRG mice. Humanized FNRG mice with serum hAlb levels of approximately 10 mg/mL were considered to exhibit >90% liver repopulation and were preferentially used for mpPHH isolation. To assess hepatocyte health in mpPHH cultures, culture supernatants were collected at the indicated time points, and hAlb levels were measured using the same ELISA assay. ELISA plates (Thermo Fisher Scientific #4424040) were coated with goat anti-human albumin polyclonal antibody (Bethyl Laboratories #A80-129A), and bound hAlb was detected using HRP-conjugated detection antibody (Bethyl Laboratories #A80-229P). Signal was developed using TMB substrate (Sigma Millipore #T0440) and stopped with sulfuric acid (Fisher Chemical # A300-500).

### FACS analysis of harvested mpPHHs

Once serum HBV DNA had plateaued, mpPHHs were harvested as described in the mpPHH section above. Four separate harvests were performed on different days with the dpi noted in Supplementary Table 6. An NT and a *ZCCHC14* KO mouse were included in each harvest. Cells were counted (Countess) and if the total number of cells exceeded 25 million, they were split into equal aliquots of <25 million cells for processing.

Cells were fixed using Cytofix (Thermo Fisher Scientific #BDB554714) using 100 µL of CytoFix solution per 1 million mpPHHs. Cells were incubated at 4°C for 15 min, protected from light. After incubation, cells were washed three times in 10 mLs of Cytoperm wash buffer (Thermo Fisher Scientific #BDB554723). Fixed and permeabilized cells were then stained with the following primary antibodies depending on the FACS analysis: Anti-mouse CD81 PE (1:40, BD #559159); Anti-human CD81 APC (1:40, BD #561958); Anti-human HLA A, B, C APC (1:40, Biolegend #311410); HBsAg Ab20 (1:200)^50^; HBcAg (1:100, Cell Marque #216-16-ASR). Primary antibody staining was performed in at most 2.5 mLs of CytoPerm wash buffer (for 25 million cells) for 1 h at room temperature under low agitation. After primary antibody staining, cells were then washed three times in 10 mLs of Cytoperm wash buffer. For HBcAg and HBsAg detection, cells were then stained with secondary antibodies: Goat Alexa Fluor 647 anti-human (1:1,000, Thermo Fisher Scientific #A-21445); Mouse anti-rabbit PE (1:100, Antigenix America #GRB10070). Staining was performed for 30 min at room temperature under low agitation and protected from light. After secondary antibody incubation, cells were washed 3 times with CytoPerm wash buffer, then resuspended in 1× DPBS supplemented with 0.5% BSA or FBS. Cells were strained through a 100 µM cell strainer, counted prior to FACS analysis, and diluted to approximately 5 million cells/mL.

For all harvests, a small aliquot of cells was processed without antibody (unstained), with no primary antibody, and each antigen-fluorophore combination individually. Cells were kept on ice, protected from light, until FACS was performed. Data were acquired on either a LSRII flow cytometer (BD Biosciences) or FACS Aria (BD Biosciences) and analyzed with FlowJo software (Tree Star).

### Data analysis and availability

All experiments were performed with at least three biological replicates unless otherwise indicated. Figure legends specify whether the symbols are individual replicates or mean values. Mean and errors bar details are described in the figure legends. Statistical analysis was performed in GraphPad Prism 10 software unless otherwise noted. All data will be made publicly available upon publication.

## Supplementary Materials

**Supplementary Fig. 1 |** Genome-wide CRISPR-Cas9 KO screens for HBV and comparisons to similar published studies

**Supplementary Fig. 2 |** Prioritization of candidate HBV pro- and anti-viral host factors and orthogonal validation of *EXOC1* in HepDE19 cells

**Supplementary Fig. 3 |** Transcriptomic changes induced by *EXOC1* depletion during HBV infection

**Supplementary Fig. 4 |** Mouse humanization and additional FACS analysis of mpPHHs isolated following HBV infection

**Supplementary Table 1:** HBsAg genome-wide screen, significant genes

**Supplementary Table 2:** HBsAg genome-wide screen, gene set enrichment analysis

**Supplementary Table 3:** Screen comparison, Venn diagram gene lists

**Supplementary Table 4:** CRISPR-Cas9 sgRNA and siRNA sequences

**Supplementary Table 5:** Screen 72 genes follow-up

**Supplementary Table 6:** Results for proviral *in vivo* genes

