## Supplementary Fig. for "Orthogonal CRISPR screens and human liver chimeric mice identify hepatitis B virus host factors"

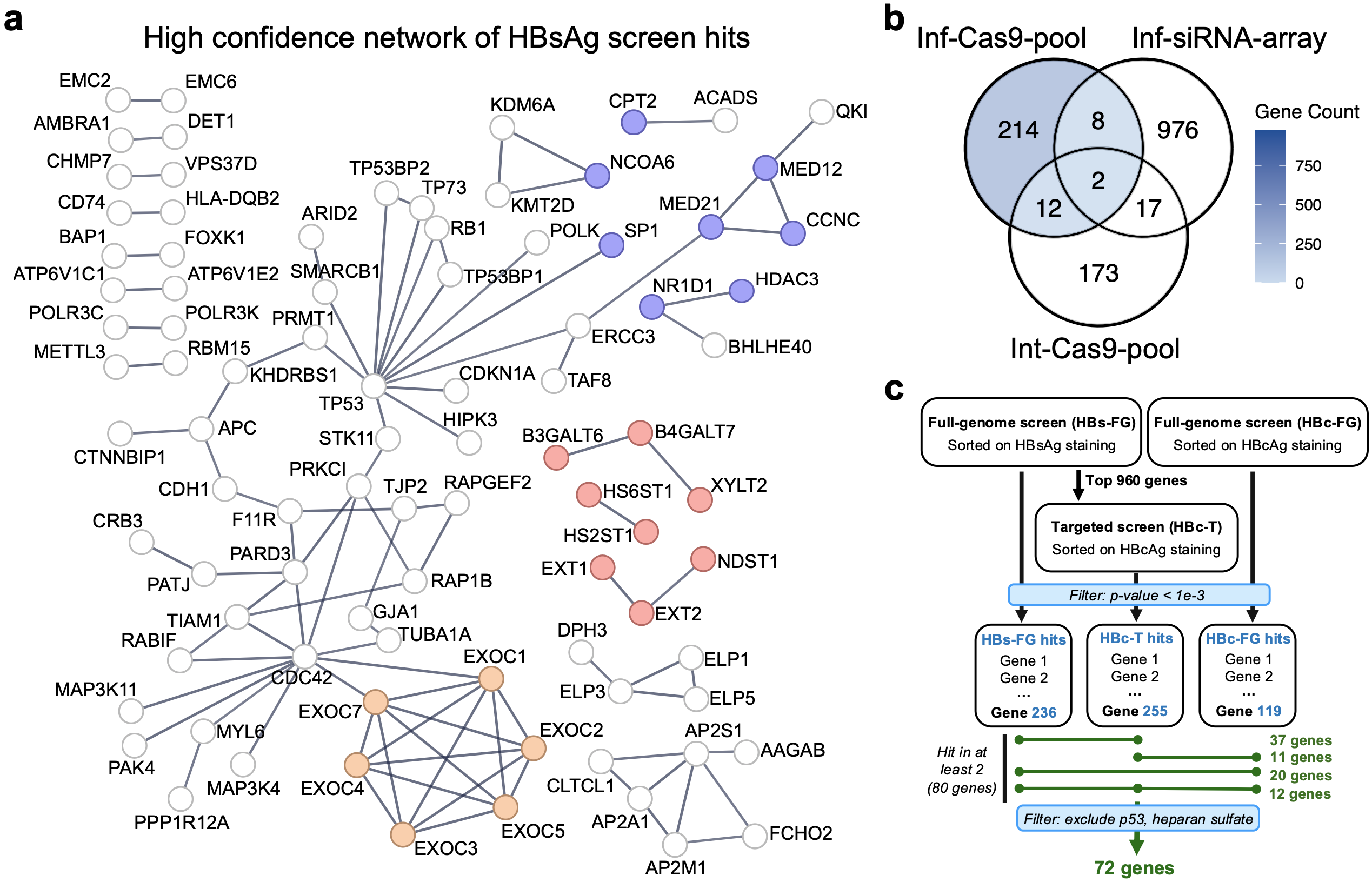

**Supplementary Fig. 1 | Genome-wide CRISPR-Cas9 KO screens for HBV and comparisons to similar published studies. a** Functional network of the significant genes from the HBsAg genome-wide CRISPR-Cas9 KO screen (genes in database = 229/236) using the STRING database^48^. Only high-confidence (score ≥ 0.9) interactions and networks of two or more genes are included. Genes contained in the enriched gene sets highlighted in Fig. 1 are colored. **b** Venn diagram of the genes passing the significance thresholds comparing the pooled CRISPR-Cas9 KO screen with HBV infection performed in this study (Inf-Cas9-pool), pooled CRISPR-Cas9 KO screen with integrated HBV system (Int-Cas9-pool)^18^, and arrayed siRNA screen with HBV infection (Inf-siRNA-array)^20^. Regions of the Venn diagram related to the significant genes in this study are shown in blue. Blue color is scaled by the number of genes in the section. See Supplementary Table 3 for the gene names. **c** Schematic of the analysis workflow for selecting the 72 genes for follow-up studies.

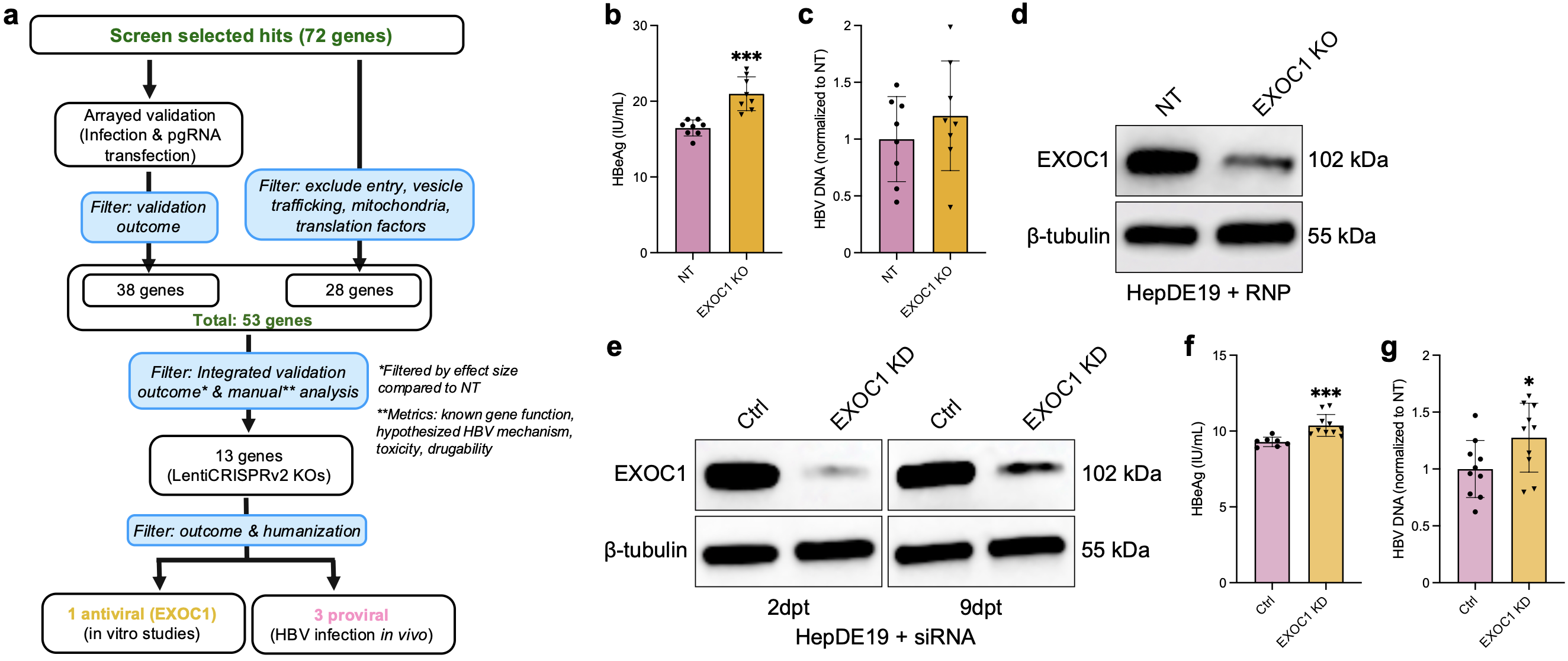

**Supplementary Fig. 2 | Prioritization of candidate HBV pro- and anti-viral host factors and orthogonal validation of *EXOC1* in HepDE19 cells.** **a** Schematic of the workflow used to prioritize candidate HBV host factors for follow-up studies. The 72 screen-identified genes were evaluated using arrayed validation assays and a literature-informed analysis. Integration of both approaches generated a list of 53 candidate genes, from which 13 genes were selected for generation of stable knockout cell lines and further validation, ultimately identifying *EXOC1* as an antiviral host factor and three proviral host factors studied in a humanized mouse model. **b,c** Validation of *EXOC1* targeting with recombinant Cas9 and synthetic sgRNAs in HepDE19 cells. NT and *EXOC1* KO cells were generated and maintained without tetracycline for at least 2 weeks. Then, cells were seeded, and 9 days post-seeding, HBV readouts were measured. HBeAg **(b)** and extracellular HBV DNA **(c)** levels were quantified. **d** Western blot analysis of EXOC1 protein levels in NT and *EXOC1* KO HepDE19 cells. β-tubulin was used as a loading control. **e** Western blot analysis of EXOC1 protein levels in HepDE19 cells transfected with control (Ctrl) or *EXOC1*-targeting siRNAs. Cells were harvested at 2 days post-transfection (dpt) and 9 dpt. β-tubulin was used as a loading control. **f-g** HBV replication readouts in HepDE19 cells following siRNA-mediated *EXOC1* knockdown (KD). Secreted HBeAg **(f)** and extracellular HBV DNA **(g)** levels were quantified following *EXOC1* KD at 9 dpt.

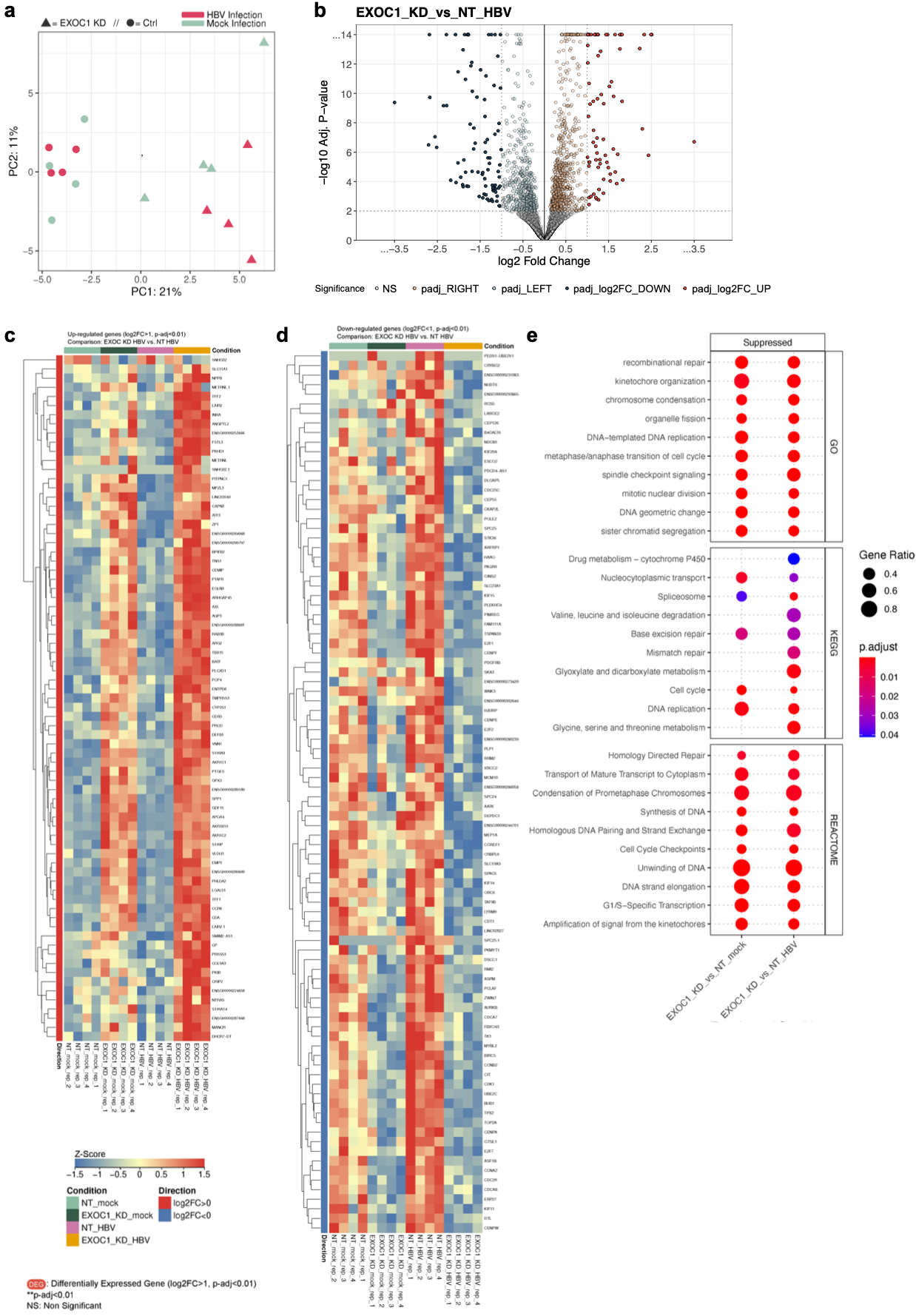

**Supplementary Fig. 3 | Transcriptomic changes induced by *EXOC1* depletion during HBV infection.** **a** Principal component analysis (PCA) of RNA-sequencing (RNA-seq) samples from NT control and *EXOC1* KD HepG2-NTCP cells in the presence and absence of HBV infection. Each point represents an independent biological replicate. Samples primarily segregated according to *EXOC1* depletion status rather than HBV infection. **b** Volcano plot showing differential gene expression between *EXOC1* KD and control HBV-infected cells. Significantly upregulated and downregulated genes were identified using an adjusted p-value cutoff of 0.01 and an absolute log_2_ fold-change (log_2_FC) threshold of 1. **c** Heatmap of significantly upregulated genes (log_2_FC > 1, adjusted *P* < 0.01) in *EXOC1* KD HBV-infected cells relative to control HBV-infected cells. Gene expression values are displayed as row-scaled z-scores. **d** Heatmap of significantly downregulated genes (log_2_FC < -1, adjusted *P* < 0.01) in *EXOC1* KD HBV-infected cells relative to control HBV-infected cells. Gene expression values are displayed as row-scaled z-scores. **e** Gene set enrichment analysis (GSEA) of pathways suppressed following *EXOC1* KD in HBV-infected cells. The most enriched downregulated pathways from Gene Ontology (GO), KEGG, and Reactome databases are shown. Dot size represents gene ratio, and color indicates adjusted *P*. Suppressed pathways were predominantly associated with cell-cycle progression, mitotic regulation, chromosome segregation, spindle organization, and DNA replication. The data represent four independent biological replicates per condition.

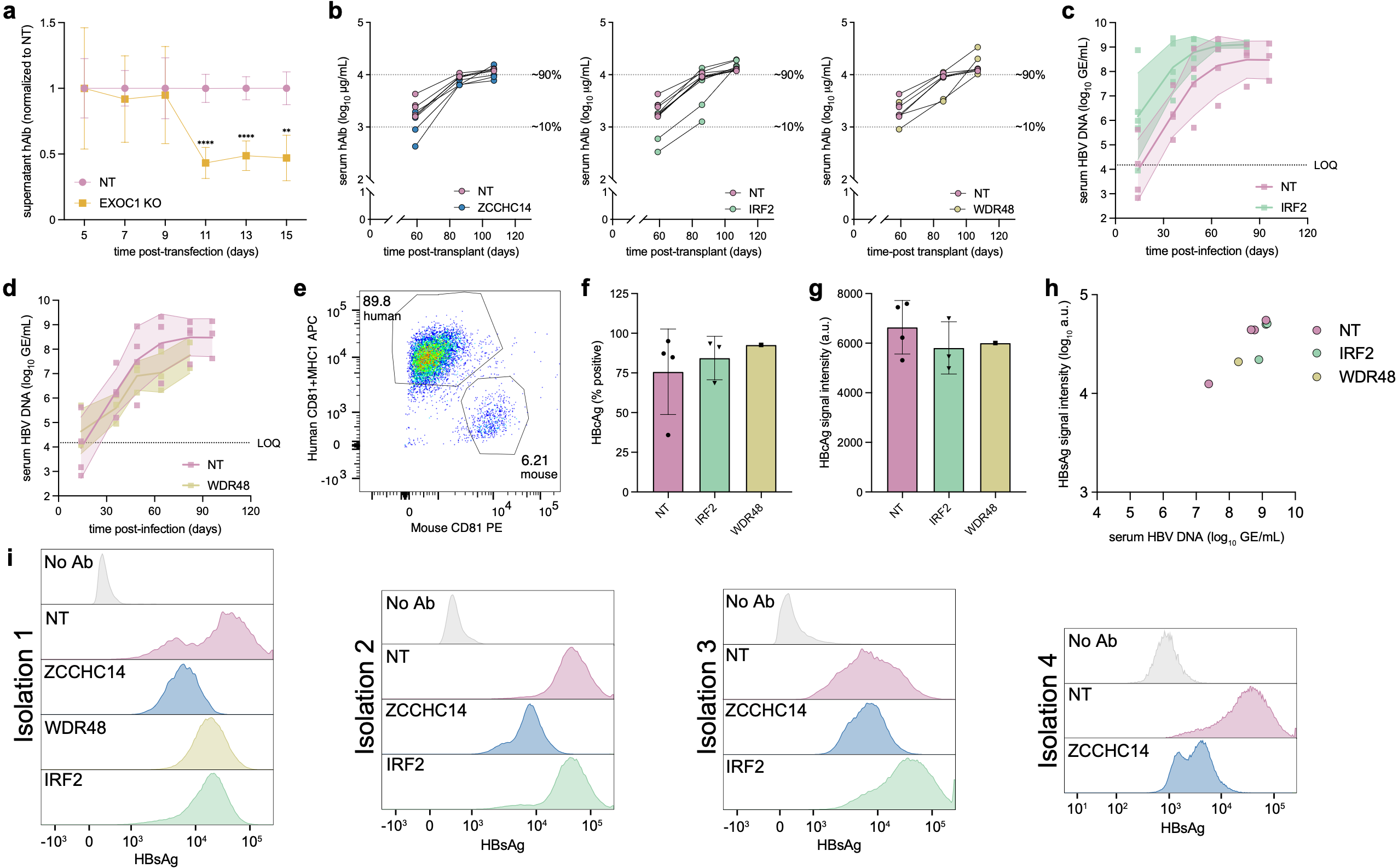

**Supplementary Fig. 4 | Mouse humanization and additional FACS analysis of mpPHHs isolated following HBV infection. a** Longitudinal analysis of hepatocyte function following *EXOC1* depletion. Secreted human albumin (hAlb) levels in culture supernatants were measured every 2 days following CRISPR-Cas9-mediated *EXOC1* KO in primary human hepatocytes. hAlb levels were normalized to day 5 for each biological replicate and subsequently expressed relative to the corresponding NT control at each time point. Data are presented as mean ± SEM of n = 6 replicates. Statistical significance was assessed using a two-way repeated-measures ANOVA with Šídák's multiple-comparisons test. ***P* < 0.01; *****P* < 0.0001. **b** Serial serum human albumin levels of FNRG mice following transplantation of mpPHHs with NT or gene-targeted sgRNAs delivered by lentiviral transduction. Each plot is an individual gene KO group with the same NT mice shown in all plots for comparison. Each symbol is an individual mouse. Mice per group: n = 6 (NT, *IRF2* KO), n = 4 (*ZCCHC14* KO), n = 3 (*WDR48* KO). **c,d** Serial serum HBV DNA levels following humanization of FNRG mice shown in **(b)** of NT mice and *IRF2* KO mice **(c)** and *WDR48* mice **(d)**. Dark line represents the mean of all mice per group; shading represents S.D.; each symbol represents an individual mouse and time point. Mice per group: n = 6 (NT), n = 5 (*IRF2* KO), n = 3 (*WDR48* KO). **e** FACS analysis of isolated NT sgRNA-transduced mpPHHs following engraftment and HBV infection. Primary antibodies were mouse CD81 (x-axis) and a pool of human CD81 and MHC-1 (y-axis) to measure the relative proportions of mouse and human cells in the population following mpPHH isolation (gates and frequencies shown as a percentage). **f** Percent of HBcAg-positive isolated mpPHHs measured by FACS. **g** HBcAg signal of HBcAg-positive isolated mpPHHs measured by FACS. **h** Serum HBV DNA (x-axis) is plotted against mean HBsAg signal intensity (y-axis). Each symbol represents an individual mouse; all mice from all isolations are included. **i** Histograms of cell-associated HBsAg signal measured by FACS. Same mice shown in Fig. 6 are reproduced here for comparison; see isolation 2. Each histogram represents data from an individual mouse. For **f-h,** the same NT mice shown in Fig. 6 are reproduced here for comparison. Each symbol represents an individual mouse. Error bars, S.D. For **f-i**, one million isolated mpPHHs from the HBV-infected NT mouse, including in each isolation, were used for a no-primary-antibody control to establish the HBsAg- and HBcAg-negative gates.
